# Harnessing tumor immune ecosystem dynamics to personalize radiotherapy

**DOI:** 10.1101/2020.02.11.944512

**Authors:** G. Daniel Grass, Juan C.L. Alfonso, Eric Welsh, Kamran A. Ahmed, Jamie K. Teer, Louis B. Harrison, John L. Cleveland, James J. Mulé, Steven A. Eschrich, Heiko Enderling, Javier F. Torres-Roca

## Abstract

Radiotherapy is a pillar of cancer care and augments the response to immunotherapies. However, little is known regarding the relationships between the tumor immune ecosystem (TIES) and intrinsic radiosensitivity, and a pressing question in oncology is how to optimize radiotherapy to improve patient responses to immune therapies. To address this challenge, we profiled over 10,000 primary tumors for their metrics of radiosensitivity and immune cell infiltrate (ICI), and applied a new integrated in silico model that mimics the dynamic relationships between tumor growth, ICI flux and the response to radiation. We then validated this model with a separate cohort of 59 lung cancer patients treated with radiotherapy. These analyses explain radiation response based on its effect on the TIES and quantifies the likelihood that radiation can promote a shift to anti-tumor immunity. Dynamic modeling of the relationship between tumor radiosensitivity and the TIES may provide opportunity to personalize combined radiation and immunotherapy approaches.

## Introduction

There is an appreciable spectrum of in vitro sensitivity to ionizing radiation across various cancer cell types that is regulated by the underlying molecular repertoire^1^ and the capacity to utilize available nutrients.^2^ However, in a tumor comprised of diverse cellular architecture and evolving ecosystems, the response to radiation is much more complex.^3^ Despite acknowledged variations in radiosensitivity, the field of radiation oncology currently does not individualize radiation dose prescription based on the intrinsic biology of a patient’s tumor.

To better understand the diversity of radiosensitivity in cancer and to identify conserved radiation response modifiers, we previously modeled the radiation response of 48 genetically annotated human cancer cell lines. By integrating the basal transcriptome, *TP53* and *RAS* isoform mutational status, tissue of origin, and clonogenic survival following a radiation dose of 2 Gy (SF_2_), we identified an interaction network of 474 genes that includes regulators of DNA damage repair (DDR) (e.g. *ATM*, *XRCC6*), oxidative stress (e.g. *PRDX1*, *TXN*), as well as ten dominant signaling hubs: *AR*, *JUN*, *STAT1*, *PKCB*, *RELA*, *ABL1*, *SUMO1*, *PAK2*, *HDAC1* and *IRF1.*^4^ From this network we derived the radiosensitivity index (RSI) by training a multi-gene algorithm to predict SF_2_ in the 48 cells lines. Notably, the radiation response signature is agnostic to cancer type and has been independently validated as a marker of radioresponsiveness across multiple human tumor types,^5–12^ and has recently been proposed as a measurable tumor feature to guide prescription of radiotherapy dose in patients.^13^

The clinical utility of harnessing a patient’s own immune system has emerged as a new pillar of tumor therapy, and as such a diverse range of tactics are being investigated to modulate the immune response, including the use of radiotherapy.^14, 15^ Indeed, emerging pre-clinical work has shown that irradiated tumors act as adjuvants for the immune system, by facilitating local tumor control and by provoking regression of tumor deposits at distant sites via the abscopal effect.^16^ Although synergism has been identified between radiotherapy and the immune system, it is not clear how to optimally integrate radiotherapy in the era of immune-modulating agents. With over three hundred ongoing clinical trials to date combining radiotherapy and immunotherapy, this is a central question in clinical oncology.

Classic radiobiology defines the radiation response as a cellular cytotoxic model based on the 5 Rs (DNA Repair, Repopulation, Reassortment, Reoxygenation and Radiosensitivity).^17^ However, the clinical response to radiotherapy occurs in the evolutionary context of complex tumor ecosystems that are comprised of multiple interacting cellular compartments. Using this dynamic system as a framework, we interrogated the relationship between the tumor-immune ecosystem (TIES) and cellular radiosensitivity in solid tumors to inform how one could harness the TIES to optimize delivery of radiotherapy.

To address this need, we initially performed a systematic multi-tier analysis of a cohort of 10,469 transcriptionally profiled, prospectively collected tumor samples representing 31 tumor types. These analyses revealed highly complex relationships between radiophenotype and the TIES that are diverse within and across tumor types. To assess how the interaction of the TIES might influence the response of tumors to radiation, we developed an *in silico* three-dimensional agent-based model that explored the dynamic interplay between tumor cells and the influx/efflux of immune cell populations during tumor development and following radiation. In this model, the TIES is a juxtaposition of two balancing phenotypes, anti-tumor versus pro-tumor, which is based on the proportion of suppressor and effector immune cell infiltrate (ICI) composition of the TIES. Finally, we develop a metric of individual radiation immune sensitivity (iRIS), which explains radiation response based on its effect on the TIES and quantifies the likelihood that radiation can shift the TIES to promote anti-tumor immunity.

## Results

### Profiling defines a spectrum of intrinsic radiation sensitivity within and across tumor types

The ‘omics’ era offers many insights into tumor biology and provides a means to classify tumor types based on certain attributes. In this regard, multidimensional reduction analysis of transcription data from 10,469 non-metastatic, primary macrodissected tumors by the t-distributed stochastic neighbor embedding (t-SNE) method demonstrates distinct clustering of tumor types (Extended Data Fig. 1).

**Fig. 1.**
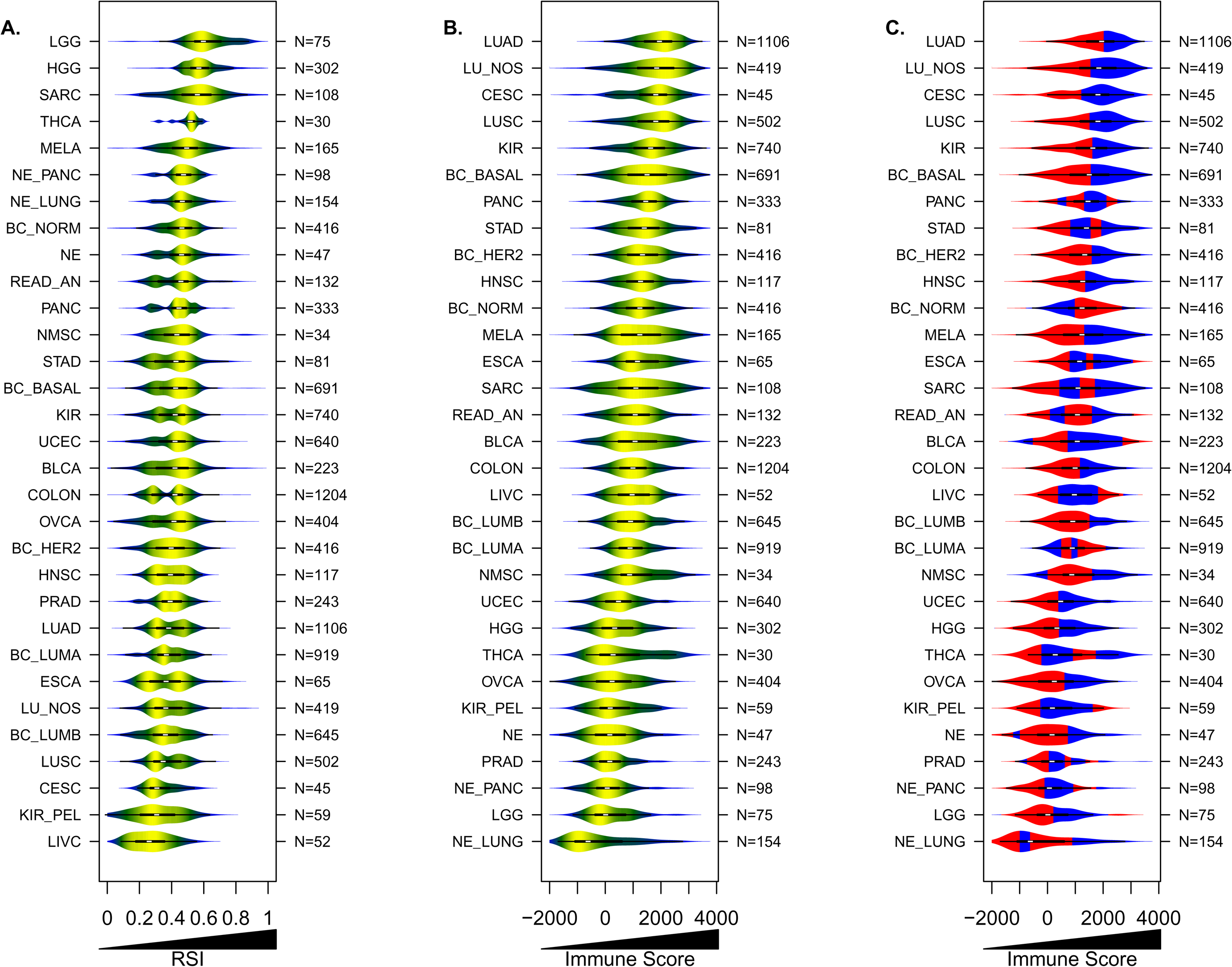
Variances and similarities in radiosensitivity and ICI presence among tumors. Violin plots depicting distribution of (**a**) RSI values and (**b**) ESTIMATE-derived immune scores across 10,469 primary tumor samples, representing 31 tumor types. **c**, Integration of RSI and the immune score. Blue (radiosensitive) and red (radioresistant) determined by the median RSI value within each tumor type.

In addition to broadly categorizing tumor types, profiling efforts have provided a basis for targeted therapy (e.g., kinase inhibitors) selection for several malignancies.^18^ In contrast, the specific radiation dose and fractionation scheme delivered to a patient’s tumor is not currently informed by tumor biology, but is rather guided by clinicopathologic features and decades of empirically derived tolerances of surrounding normal tissues. Further, although substantial experimental and clinical evidence suggest tumors in the same anatomic location and of similar histological subtype can vary widely in their response to radiation, a uniform dosing scheme is generally employed in clinical practice.

To evaluate the radiosensitivity of human tumors, we estimated intrinsic radiation sensitivity using the radiosensitivity index (RSI)^9–11^. Notably, a spectrum of RSI values within and across the 31 specified tumor types is evident (Fig. 1a). For example, concordant with clinical experience, the most radioresistant (highest median RSI) tumors are gliomas and sarcomas, whereas tumors of the liver, renal pelvis and cervix are the most radiosensitive (lowest median RSI). Stratification of breast cancers into genomically-derived subtypes^19, 20^ revealed normal-like breast tumors are the most radioresistant, whereas luminal A and B types are the least (*P* < 0.001). In contrast, no differences in the median RSI are noted between histologic subtypes of lung cancer (*P* = 0.096). Evaluation of interquartile range (IQR) ratios for RSI reveal that renal pelvis, liver, gastrointestinal or genitourinary originating tumors have the greatest dispersion, whereas thyroid and neuroendocrine tumors, and the more radioresistant gliomas, sarcomas and pancreatic tumors are more uniformly distributed (**Supplemental Table 1**). Finally as a whole, the RSI distribution among all tumor types is not unimodal based on Hartigan’s dip test statistic,^21^ where several tumor types have non-unimodal distributions in RSI, including kidney, large bowel, rectum-anus, prostate, esophagus, pancreas, stomach, and lung and breast subtypes (Extended Data Fig. 2).

**Fig. 2.**
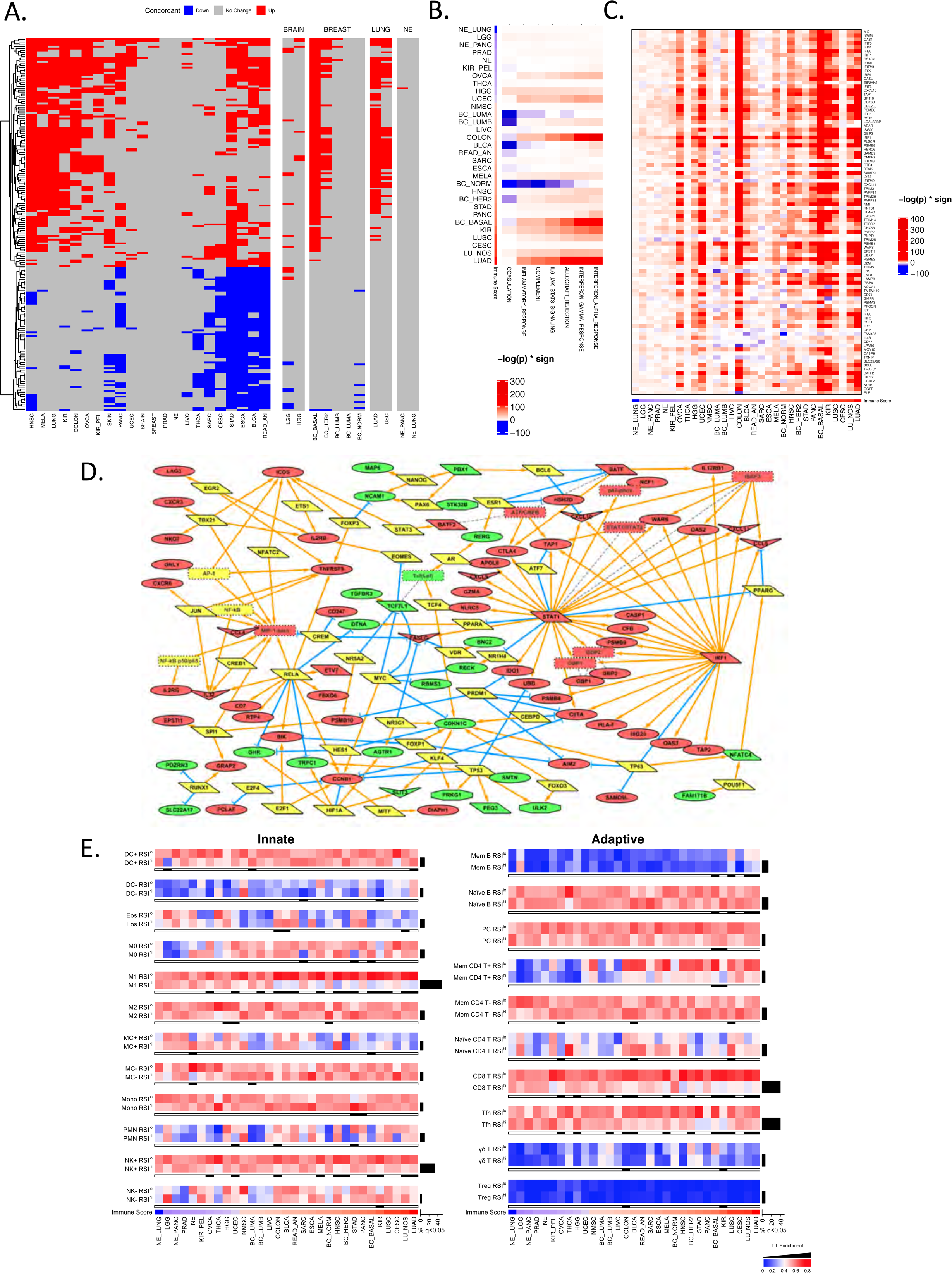
IFN signaling pathways influence the immune portraits of intrinsic tumor radiosensitivity. a) Heatmap of differentially expressed probesets (n=197, representing 146 unique genes) between radiosensitive (RSI^lo^)and radioresistant (RSI^hi^) tumors across six or more of the 31 tumor types, which are concordant in direction of expression (up, red; down, blue; no change, gray). See Supplemental Table 3 for list of probesets. **b)** RSI^lo^ versus RSI^hi^ tumors were compared with single sample gene set enrichment analysis (ssGSEA) using the MSigDB hallmark pathway genes related to immune signaling. Heatmap intensity depicts the -log p-value of the comparison test multiplied by the directionality of expression difference. **c)** Individual genes (n=97) of the MSigDB IFNα hallmark pathway. Tumor types are ordered by the ESIMATE-derived immune score and heatmap intensity depicts the -log p-value of the comparison test multiplied by the directionality of expression difference of RSI^lo^ versus RSI^hi^ tumors. **d)** Differentially expressed probesets (146 genes) from Fig. 2a were used as seeds for network generation (see Methods). Genes upregulated (red), downregulated (green) in RSI^lo^ versus RSI^hi^ tumors are depicted. Yellow nodes indicate bridging genes. **e)** Enrichment of different ICIs between RSI^lo^ and RSI^hi^ tumors. Tumor types are ordered by the ESTIMATE-derived immune scores and normalized CIBERSORT-derived ICI estimates are compared within a given tumor type by dichotomizing at the median RSI value to identify RSI^lo^ and RSI^hi^ tumors. ICIs involved in the innate response (*left panel*) and ICIs involved in the adaptive response (*right panel*). Black bars, proportion of tumor types for given ICI enrichment comparison between RSI^lo^ and RSI^hi^ tumors; false-discovery rate < 0.05.

### Increased ICI abundance connotes radiosensitivity across most tumor types

Diverse cell types, including innate and adaptive immune cells and stromal cells, influence the dynamics and radiophenotype of tumors.^22^ The ESTIMATE algorithm^23^ was employed to infer the fraction of each respective cell type and to approximate tumor purity (i.e., malignant cell burden) within each tumor. The distributions of ESTIMATE-derived stromal, immune and tumor purity scores were visualized for all tumors samples analyzed (Extended Data Fig. 3).

**Fig. 3.**
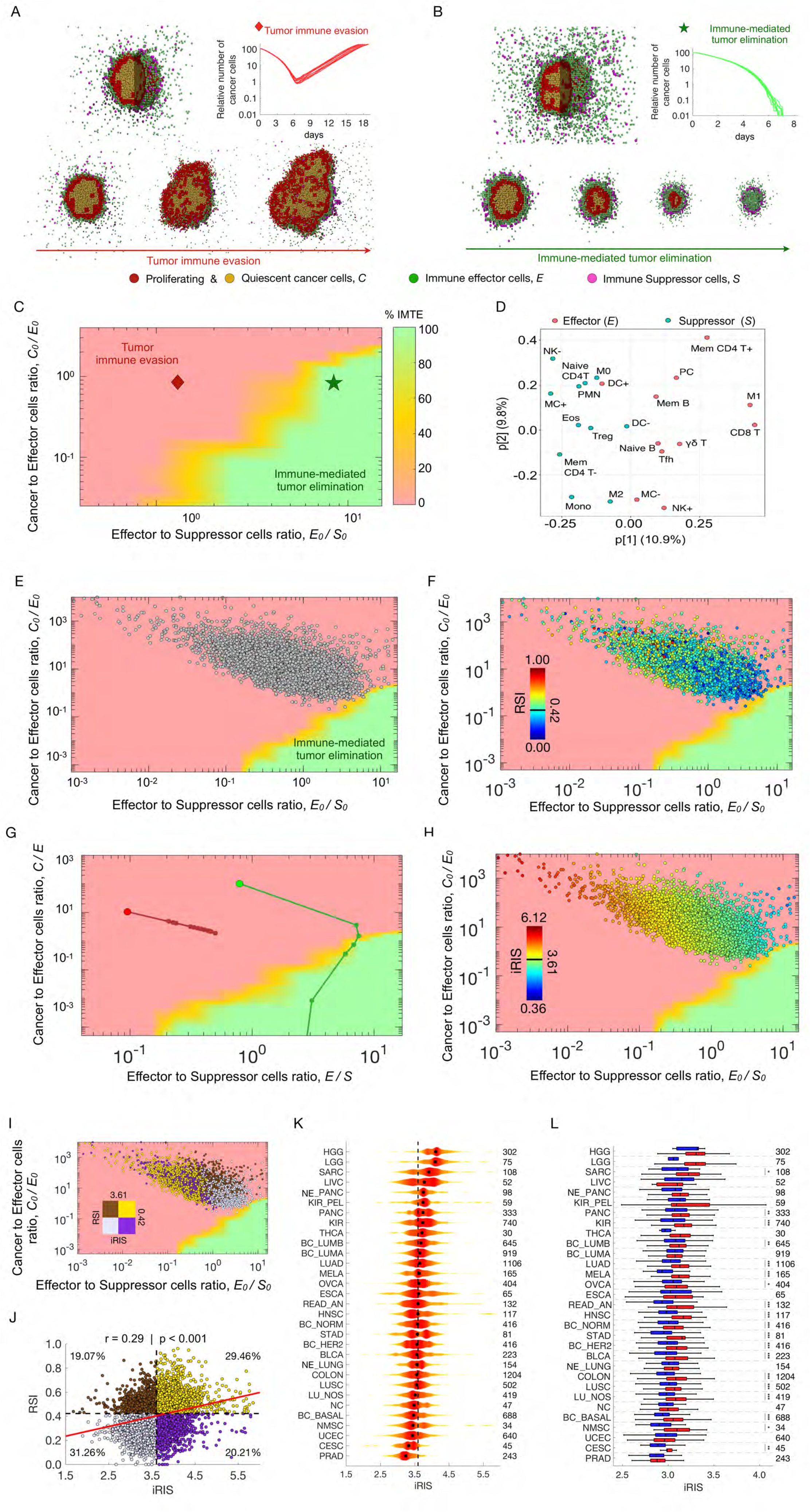
Integrative *in silico* modeling defines the tumor immune ecosystem (TIES) and the individual radiation immune sensitivity (iRIS) score. Biologically-defined mechanistic rules guiding tumor cell and ICI interactions in the 3D agent based model (see Methods). A fixed starting malignant cell burden and varying proportions of effector or suppressor ICIs were inputted into the model and simulations resulted in **a)** tumor immune evasion or **b)** immune-mediated tumor elimination. Different populations of cells in the agent-based model are depicted as follows: red (proliferating tumor cells), tan (quiescent tumor cells), green (effector ICIs) and purple (suppressor ICIs). **c)** Representation of probability of immune-mediated tumor elimination (IMTE) based on individual tumor immune ecosystems (TIES) derived from Fig 2a,b. y-axis: ratio of malignant cell burden/effector ICI (*C_0_/E_0_)*; x-axis: ratio of effector/suppressor ICI (*E_0_/S_0_).* Red region, TIES characterized by tumor immune evasion; green region, TIES representing immune-mediated tumor elimination. **d)** Loading of principal component analysis (PCA) of the CIBERSORT-derived ICI composition across 10,469 tumors stratified as anti-tumor (*E*; effector) or pro-tumor (*S*; suppressor). **e)** The cellular composition (normalized ICI counts and malignant cell burden estimated by CIBERSORT and ESTIMATE, respectively) for each of the 10,469 tumors was derived and plotted onto the TIES map; all tumors localized in regions with a TIES leading to tumor immune evasion. **f)** Visualization of continuous RSI values for each tumor on the TIES map reveals more radiosensitive (RSI^lo^ based on population median) tumors are in closer proximity to TIESs which promote immune-mediated tumor elimination. **g)** Modeled trajectories of TIES composition evolution of a given pre-treatment TIES following radiation treatment. Each closed circle on the trajectory represents a radiation dose of 2 Gy/day. This demonstrates that radiation causes shifts in the TIES, which result in different trajectories despite being equivalent closest Euclidean distances from the IMTE region (green). **h)** The individual radiation immune sensitivity (iRIS) score for each tumor, which describes the specific position in the TIES map with respect to the shortest Euclidean distance from the IMTE region, penalized by the absolute number of suppressor ICIs. A lower iRIS value represents a more radiation responsive TIES. **i)** Stratification of tumors by the median RSI and iRIS values distinguishes specific TIES and probability of immune-mediated tumor elimination. **j)** Scatter plot with correlation of RSI and iRIS across all tumor types. Pearson’s r = 0.29, *P* < 0.001. Percentages depict proportion of specific TIES in quadrants defined by the median RSI and iRIS values. **k)** Violin plots depicting distribution of iRIS values, highlighting heterogeneity within and across tumor types. **l)** Boxplots comparing iRIS distributions between RSI^lo^ and RSI^hi^ tumors (Mann Whitney U test; * *P* < 0.05, ** *P* < 0.01, *P* < 0.001). Demonstrates that most radiosensitive tumors are associated with a lower iRIS.

Similar to the variance observed with the RSI, ESTIMATE-derived approximations of non-tumor cell infiltrates identified a wide range of ICI (Fig. 1b) and stromal (Extended Data Fig. 4) cells within and across tumor types. Tumor types with clinical responsiveness to immune checkpoint blockade therapies^24^, such as lung, kidney and melanoma, or those largely driven by viral infection (e.g., cervix), have the highest median immune scores, which reflects ICI abundance, especially tumor infiltrating lymphocytes (TIL). Additionally, consistent with prior observations,^25^ the breast basal subtype has the highest presence of ICI. Tumors with the lowest median immune scores are gliomas, prostate cancer and neuroendocrine tumors.

**Fig. 4.**
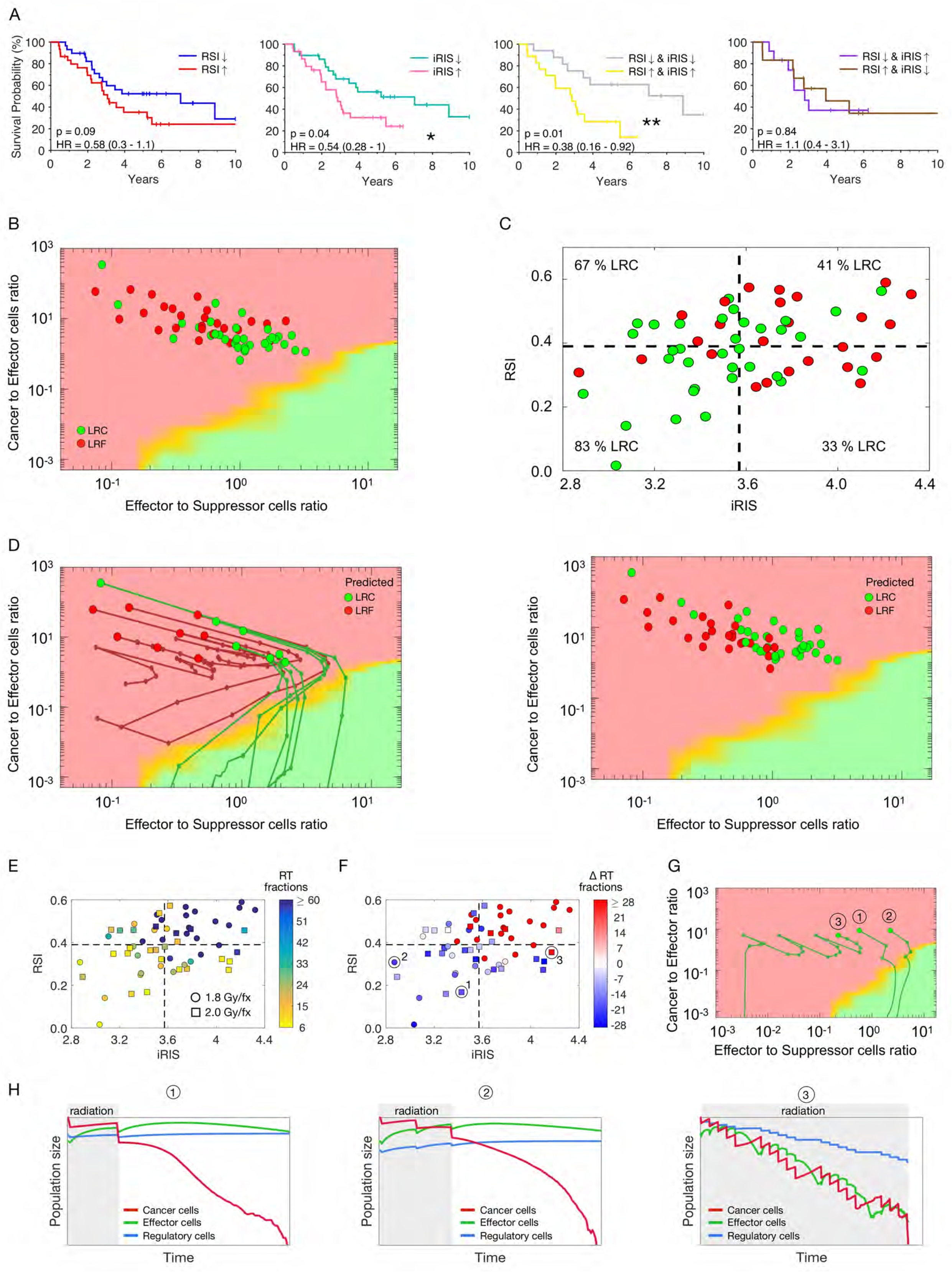
*In silico* modeling optimizes radiation dose delivery based on the pretreatment TIES in non-small cell lung cancer patients. a) An independent cohort of 59 non-small cell lung cancer (NSCLC) patients treated with postoperative radiation at various doses (range: 42-70 Gy) were analyzed for locoregional control (LRC), failure (LRF) or overall survival (OS). **a)** Kaplan-Meier estimates for OS demonstrate patients with tumors classified as RSI^lo^ vs RSI^hi^ have a trend for improved OS (hazard ratio: 0.58, range: 0.3-1.1; *P*-value = 0.09), whereas iRIS^lo^ vs iRIS^hi^ tumors (hazard ratio: 0.54, range: 0.28-1.0; *P*-value = 0.04) have improved OS. Patient tumors with dual RSI^lo^/iRIS^lo^ phenotypes have improved OS (hazard ratio: 0.38, range: 0.16-0.92; *P*-value = 0.01) compared to those with RSI^hi^/iRIS^hi^, though a RSI^lo^ phenotype with an unfavorable TIES (iRIS^hi^) or vice versa, cannot compensate for the other (hazard ratio: 1.1, range: 0.4-3.1; *P*-value = 0.84). **b)** The cellular composition (ICIs and malignant cell burden) of each patient tumor was plotted onto the TIES map and actual clinical outcomes of LRC (green) and LRF (red) were evaluated with respect to their given TIES. **c)** Separation of tumors by median RSI and iRIS demonstrates that 83% of RSI^lo^/iRIS^lo^ tumors achieve LRC compared to 67%, 41% and 33% in RSI^hi^/iRIS^lo^, RSI^hi^/iRIS^hi^, and RSI^lo^/iRIS^hi^, respectively. **d)** (*left* panel) Representation of a subset (n=15) of analyzed tumors each undergoing *in silico* modeling according to methods of Fig. 3a,b, with each tumor receiving radiation treatment according to the actual delivered treatment protocol (dose per fraction and total dose). The projected shift in the individual TIES following radiation is depicted by the trajectory; the simulation was continued until complete response (i.e., no tumor cells alive) or treatment failure (i.e. returned to pretreatment tumor burden). The space between each point on a trajectory indicates the change in TIES composition after delivery of a single fraction of radiation. Each individual tumor was grouped according to predicted outcome; LRC (green) or LRF (red). (*Right* panel) All 59 tumors were plotted onto the TIES map and categorized according to predicted outcome. **e)** Each patient tumor underwent *in silico* modeling with a repeating single 1.8 Gy (circle) or 2 Gy (square) per fraction regimen (based on actual delivered treatment regimens) until complete response was achieved or delivery of at least 60 fractions. After each individual dose the final TIES trajectory was analyzed for response. Plotted coordinates demonstrate the amount of fractions that were necessary to achieve complete response, relative to tumor specific RSI and iRIS continuous values. Notably, several tumors reach at least 60 fractions of radiation without a complete response, whereas some achieve complete response in less than 24 fractions. **f)** Difference in actual delivered radiation fractions compared to *in silico* model predicted number of fractions needed to achieve complete response. Negative values imply a decrease in total fractions, which may provide opportunity for de-escalation of treatment, whereas positive values suggest an increased number of fractions may be needed for optimal tumor control. **g)** Three selected patient tumors (circled tumors in Fig 4f) with predicted TIES shifts following simulated radiation illustrate three separate scenarios: **h)** Tumor 1) TIES promotes accelerated tumor eradication and potential for de-escalation of dose, Tumor 2) TIES promotes tumor eradication, but requires more radiation priming, or Tumor 3) TIES is immunosuppressive and only results in tumor kill after several rounds of cytotoxic radiation without assistance from the host immune system. Gray shaded area indicates course of fractionated radiation over a given time.

Due to burgeoning data supporting an interplay between radiotherapy and the immune system^14–16^, as well as data suggesting the RSI is partially associated with a 12-chemokine gene signature across several tumor types^26^, we tested if there was a relationship between intrinsic tumor radiosensitivity and the presence of ICI. Among all tumors, there is a weak negative correlation between the RSI and immune score (r = −0.28; *P* <0.001), yet principal component analysis (PCA) revealed subset aggregation of radiosensitive (i.e., low RSI values) and ICI-rich tumors in the same space (Extended Data Fig. 5). This suggests radiosensitive tumors having higher ICI. Thus, we evaluated the correlation between the RSI and immune score for each tumor type, which identified moderate negative correlations for thyroid (r = −0.56), cervical (r = −0.54), melanoma (r = −0.47), head and neck (r = −0.45), and basal breast subtype (r = −0.44) tumors; the range of Pearson’s r values for all tumor types is −0.56 - +0.024 (Extended Data Fig. 6). Similarly, stromal cell presence (r = 0.03) and tumor purity (r = 0.13) are weakly related with the RSI. Finally, as previously shown^23, 27^, tumor purity is strongly associated with the immune (r = - 0.93) and stromal (r = −0.90) scores.

Next, we classified each tumor sample within a given tumor type as radiosensitive (RSI^lo^) or radioresistant (RSI^hi^) based on the median RSI value within a given tumor type. Notably, integration of the RSI and immune score revealed that most radiosensitive tumors are characterized by increased ICI abundance across most tumor types (Figure 1c). However, there were exceptions, as some tumors classified as radiosensitive had reduced ICI, while others that were radioresistant had high ICI. Overall, the data imply that for most tumor types, ICI presence connotes radiosensitivity.

### IFN signaling connotes intrinsic tumor radiosensitivity

To identify the biological discriminators of RSI^lo^ and RSI^hi^ tumors, each tumor type was evaluated for differentially expressed genes (DEGs) (**Supplemental Table 2**). Across all tumor types there were 7,184 DEGs between the RSI groups. Interestingly, prostate, neuroendocrine or non-subtyped breast tumors have two or less RSI-influenced DEGs, whereas breast tumor subtypes, except luminal variants, have greater than 150 DEGs between the RSI^lo^ and RSI^hi^ groupings. Furthermore, neuroendocrine tumors stratified as being of pancreas or lung origin unmasked additional DEGs, underscoring the contribution of the microenvironment.

We further investigated whether a conserved RSI-influenced transcriptional program is present in tumors. Notably, these analyses identified 209 unique probesets (155 genes) that are differentially expressed between RSI^lo^ and RSI^hi^ tumors across at least six tumor types, and of these, 146 genes are concordant in their direction of expression (Fig. 2a and **Supplemental Table 3**).

Given the relationships between radiosensitivity, immune signaling and the presence of ICI, single sample gene set enrichment analysis (ssGSEA) was performed with the immune-related MSigDB hallmark gene sets.^28^ Across several tumor types, ssGSEA revealed that RSI^lo^ versus RSI^hi^ tumors are enriched in pathways regulating type I (α) and II (γ) interferon (IFN) signaling (Fig. 2b). Further, RSI^lo^ tumors having the greatest enrichment of IFNα or IFNγ signaling were mostly manifest in tumor types classified as having the highest immune scores. Evaluation of the IFNα signaling module revealed that numerous genes involved in various aspects of this signaling network are upregulated in RSI^lo^ versus RSI^hi^ tumors (Fig. 2c). Similarly, genes representing six other immune-related pathways were also upregulated in RSI^lo^ tumors across various tumor types (Extended Data Fig. 7). Pathway topology analyses with the 146 conserved genes identified an expansive interacting network with STAT1, IRF1, and CCL4/MIP-1β as major upregulated nodes in RSI^lo^ tumors (Fig 2d).

### Tumor radiosensitivity correlates with select ICI composition

The ESTIMATE-derived immune score provides a generic metric of ICI presence, and does not elucidate the composition or functional state of ICIs.^23^ To address the immune repertoire and activation state of tumors, the CIBERSORT deconvolution algorithm^29^ was used to infer the presence of 22 distinct immune cell subtypes within each tumor ICI. We found that ICIs involved in the adaptive or innate immune responses were differentially enriched between RSI^lo^ (radiosensitive) and RSI^hi^ (radioresistant) tumors within each tumor type. A common pattern of ICI enrichment was seen in RSI^lo^ versus RSI^hi^ tumors for CD4+ memory T cells (Mem CD4^+^), CD8+ T cells (CD8^+^ T), follicular helper T cells (Tfh), activated natural killer (NK) cells (NK^+^) and M1-polarized macrophages (M1) (Figure 2e; q-value < 0.05). However, there is a significant heterogeneity in ICI enrichment both across and within tumor types.

Although relationships between radiosensitivity and ICI presence and composition are apparent, other variables influence both. Indeed, the mutational landscape has been shown to influence tumor immune responses^30–34^ and previous investigations have identified certain mutations that may correlate with radiosensitivity^1, 35^. To address this relationship we performed targeted sequencing of a subset of tumors (n=2,368) across all types. Notably, mutation frequency (both non-synonymous and synonymous) only weakly correlated with the RSI (r = −0.07; *P* = 0.001), immune score (r = −0.01; *P* = 0.54) and ICI composition (**Supplemental Table 4**).

### Dynamic tumor-immune ecosystem models define a novel individual radiation immune sensitivity metric

The tumor ecosystem is a dynamic and diverse network of cellular and non-cellular constituents that can either perpetuate or attenuate tumor growth. To model the dynamic interplay between tumor cells and the influx and efflux of ICI components, we generated an *in silico* 3-dimensional agent-based model guided by biologically defined rules, which incorporates varying proportions of effector and suppressor immune cell populations. The absolute numbers of the effector and suppressor ICI cells along with the cancer cell burden were used to define the tumor-immune ecosystem (TIES, see Methods). Simulation of tumor growth with various TIESs reveals two achievable outcomes, those where tumors evade immune predation (Fig. 3a) and those where tumors are eradicated by the immune system (Fig. 3b). The ratio of cancer cells to effector ICI cells (*C_0_/E_0_*, y-axis) plotted against the ratio of effector ICI cells to suppressor ICI cells (*E_0_/S_0_*, x-axis) spans the “TIES map”, where each pixel represents a unique TIES. Outcome statistics (immune-mediated tumor elimination; IMTE) for ten independent simulations for each initial TIES (*C_0_, E_0_, S_0_*) identifies TIES compositions that will lead to immune eradication of tumors or, alternatively, that will lead to tumor escape (Fig. 3c).

To validate these simulations, the TIES composition for each of the previously analyzed 10,469 tumors was constructed using the ICI counts and malignant cell burden from CIBERSORT-derived assessments and ESTIMATE data, respectively; ICIs were further grouped into effector and suppressor types based on PCA (Fig 3d). This method identifies that the untreated TIES (*C_0_, E_0_, S_0_*) of all 10,469 tumors as having a phenotype characterized by immune evasion (Fig 3e; Extended Data Fig. 8). Notably, superimposing tumor-specific RSI onto the TIES map suggests that tumors with lower RSI values cluster closer to TIES compositions characterized by immune-mediated tumor elimination (**Fig**. **3f**).

To further evaluate these findings, the effects of radiation on the tumor and the TIES composition were simulated by extending the agent-based model to incorporate radiation-induced cytotoxicity of cancer cells and ICIs using experimentally defined radiation sensitivities and ICI responses to radiation (see Methods). This model allows simulation of radiation-induced shifts in TIESs during fractionated radiotherapy for tumors with different pre-treatment TIES compositions. Notably, these simulations revealed that the TIES with more favorable *E_0_/S_0_* ratios have a higher probability of being shifted to post-radiation TIES compositions that promote immune eradication of residual tumor cells (Fig. 3g). From these findings a new individual radiation immune sensitivity (iRIS) metric was derived with

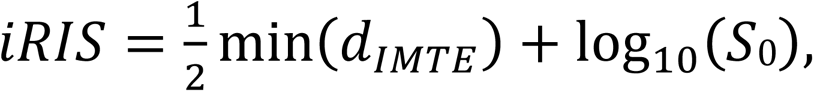

where

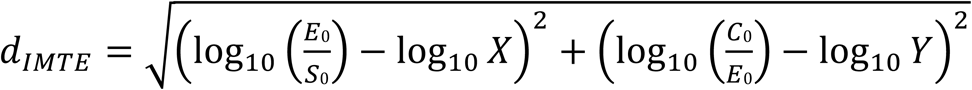

is a function of the distance of pre-radiation TIES composition to the IMTE region in the TIES map (Fig. 3h). Here the X and Y vectors are the coordinates of all TIESs in the region of the IMTE, such that min(*d_IMTE_*) denotes the trajectory with the shortest Euclidean distance from the current TIES of immune evasion to one of immune-mediated control.

Stratification into RSI^lo^ and RSI^hi^ (median RSI value for all 10,469 tumors), as well as iRIS^lo^ and iRIS^hi^ (median iRIS value) into quadrants, distinguishes TIES composition (Fig. 3i). Specifically, RSI^lo^ tumors have lower iRIS (median 3.52; SD 0.36) than RSI^hi^ (median 3.70; SD 0.42; *P* < 0.001) tumors. Similarly, iRIS^lo^ tumors have lower RSI (median 0.37; SD 0.11) than iRIS^hi^ (median RSI 0.44; SD 0.12; *P* < 0.001) tumors. Accordingly, there is a weak positive correlation between RSI and iRIS across all tumor types (r = 0.29; *P* < 0.001, Fig. 3j); the range of Pearson’s r values across tumor types is −0.02 - +0.45 (Extended Data Fig. 9). Finally, although iRIS varies across and within different tumor types (Fig. 3k), most RSI^lo^ tumor types have significantly lower iRIS (Fig. 3l).

### RSI and iRIS mutually predict radiation response in lung cancer

To validate these RSI and iRIS findings, an independent clinical cohort of 59 non-small cell lung cancer (NSCLC) patients was examined where patients treated with varying doses of post-operative radiation had overall survival (OS), locoregional control (LRC) and failure (LRF) data.

Notably, patients with RSI^lo^ tumors have a trend for improved OS (HR=0.58; *P* = 0.09), and those with iRIS^lo^ versus iRIS^hi^ tumors have improved OS (HR=0.54; *P* = 0.04; Fig. 4a). Further, as predicted, patients with dual RSI^lo^/iRIS^lo^ tumor phenotypes have superior OS versus patients with RSI^hi^/iRIS^hi^ tumors (HR=0.38 *P* = 0.01; Fig. 4a). Nonetheless, a RSI^lo^ phenotype with an unfavorable TIES (iRIS^hi^) or vice versa, does not connote improved OS (Fig. 4a); thus, both metrics mutually predict the effects of radiation on immune-mediated tumor eradication.

Plotting each patient-specific tumor and their associated outcomes onto the TIES map revealed that most tumors that achieved LRC were in closer proximity to the immune-mediated tumor elimination region (Fig. 4b). Next, to assess the ability to predict patient outcomes, the RSI and iRIS of each tumor was calculated. The total radiation dose received was not informed by RSI or iRIS, nor were RSI or iRIS correlated. Patients that achieved LRC have higher *E_0_/S_0_* (median 1.02; SD 0.74 vs median 0.51; SD 0.55, *P* < 0.001) and lower *C_0_/E_0_* ratios (median 2.62; SD 58.25 vs median 8.31; SD 18.16, *P* < 0.001). Achievement of LRC is also associated with lower iRIS (*P <* 0.01), but not RSI (Extended Data Fig. 10). Patients with favorable radiation sensitivity and TIES phenotypes (RSI^lo^/iRIS^lo^) had higher rates of LRC versus those with unfavorable radiation sensitivity and TIES phenotypes (RSI^hi^/iRIS^hi^) (83% *vs.* 41%; *P* < 0.001, Fig. 4c)

### Simulation of clinically applied radiotherapy dosing regimens provides opportunity to personalize radiation delivery based on pre-treatment TIES composition

To test if *in silico* agent-based models can inform personalized treatment, the pre-treatment TIES composition, RSI and iRIS of each NSCLC patient was derived as above. We then integrated these tumor metrics into a simulation to determine the effects of clinically applied individual radiation dosing regimens (total dose applied in 1.8-2 Gy per fractions for a total of 42-70 Gy) to predict radiation-induced shifts in TIES composition. For patients with ‘cold’ TIES (i.e., low *E_0_/S_0_* ratios) radiation fails to sufficiently perturb the TIES to promote immune-mediated tumor elimination, resulting in LRF. In contrast, in patient tumors with ‘hot’ TIES (i.e. high *E_0_/S_0_* ratios) predicted to be controlled by radiation, there are radiation-mediated trajectory shifts towards more favorable TIES compositions that support tumor elimination, either by radiation alone or by induced TIES compositions that facilitate immune-mediated eradication (Fig. 4d).

The *in silico* agent-based models were then used to predict the minimum required number of 1.8 Gy or 2 Gy fractions to eliminate patient-specific tumors. For selected RSI^lo^/iRIS^lo^ tumors, simulations revealed that as few as 6-10 fractions (10.8 −20 Gy total dose) are predicted to be sufficient to provide LRC via robust stimulation of anti-tumor immunity. In contrast, in tumors with an unfavorable RSI^hi^/iRIS^hi^ TIES compositions, simulations predict up to 60+ fractions (>120 Gy total dose) may be necessary to eradicate every single cancer cell with radiation (Fig. 4e).

Notably, of the 59 analyzed NSCLC patient-specific tumors, 6 (10%) received a radiation regimen within +/- 5 fractions of the calculated required dose by the RSI/iRIS-informed *in silico* model, whereas 30 (51%) are predicted to be candidates for radiation de-escalation, and 23 (39%) would require dose escalation by more than 5 fractions (Fig. 4f). Analysis of trajectory shifts in the TIES during radiation for three select patients (Fig. 4g) and the corresponding change in *C_0_, E_0_, S_0_* populations over time indicate two different radiation prescription purposes: 1) radiotherapy as an ICI-stimulating agent with opportunity for de-escalated doses (Fig. 4h**, left and middle panels**) or 2) radiation as a purely cytotoxic agent that has to eradicate every cancer cell without the support of the ICI (Figure 4h**, right panel**). Notably in the latter, the persistence of *S_0_* populations may propagate a microenvironment of radiation-induced, immunosuppressed ICI composition with no observable benefit to radiotherapy.

## Discussion

We believe this study represents the largest pan-cancer analysis of primary human tumors characterized as radioresistant or radiosensitive. By employing multi-tier computational analyses, several important tumor features were identified that impact efforts to personalize radiotherapy delivery and to strategically integrate radiotherapy and immunotherapies.

First, RSI, a cancer type agnostic gene signature, allowed characterization of intrinsic tumor radiosensitivity across 31 tumor types, and revealed that RSI predicts clinical experience, where ‘resistant’ (glioma, sarcoma, melanoma) and ‘sensitive’ (cervical, liver) tumors have some of the highest and lowest median RSI values, respectively. Importantly, though differences in radiosensitivity are evident between and within tumor types, the dispersion of RSI values revealed the most sensitive tumors in a given tumor type can overlap with the most resistant tumors of another type and vice versa. Further, characterization of RSI distributions by IQR ratios or dip statistics, establishes that many clinically-defined radioresistant tumors have less dispersion and unimodality, suggesting these tumor types lack variation in radioresponsiveness and that tumor control with radiotherapy alone will prove difficult. In contrast, for tumor types with greater dispersion and non-unimodal RSI distributions our data indicates there is an opportunity to personalize radiotherapy to improve the clinical response.

Radiosensitivity is influenced by complex interactions between intrinsic polygenic traits, microenvironment dynamics, utilization of nutrients and diverse cellular composites.^2, 3^ Recent studies in 533 tumor cell lines found that intrinsic radiosensitivity is interconnected with DDR and genomic stability^1^, supporting the long held tenet that radiation sensitivity is determined by the fidelity of DDR. However, tumor radiosensitivity is also governed by the other ‘hallmarks of cancer’^36^, and with the success of immunotherapies in the clinic, there is a need to deeply understand relationships between radiation and the host immune response. For this reason, we characterized the presence of ICIs across 31 tumor types and identified a wide spectrum of ICI abundance and composition across and within tumor types, confirming previous observations.^23, 37, 38^ Notably, integration of ICI presence and RSI groupings revealed that across most tumor types, increased ICI presence is associated with a radiosensitive phenotype and that certain ICI compositions are more enriched in radiosensitive versus radioresistant tumors. Similar relationships between radioresponsiveness and select features of the ICI have been identified in breast^39^, prostate^40^, and bladder^41^ cancers. Together, our studies suggest the heterogeneity in the repertoire of ICI compositions among tumor types may inform strategies that integrate radiation and immunotherapies to improve outcomes.

Recent data have suggested an interesting connection between DDR and antitumor immunity via conserved mechanisms that detect cytosolic nucleic acids to combat foreign pathogens.^42^ For example, cyclic GMP-AMP synthase (cGAS)/stimulator of IFN genes (STING) signaling, major regulators of type I IFN production^43^, influence the radiation response.^44^ Further, type I IFNs are known to stimulate both the innate and adaptive arms of the immune response^45^ and are also essential for radiation efficacy.^46, 47^ In accord with these studies, our analyses revealed that RSI^lo^ versus RSI^hi^ tumors are indeed highly enriched in inflammatory signaling, including type I IFNs. Further, although the signaling effectors *MB21D1*/cGAS and *TMEM173*/STING failed to meet our statistical criteria for concordant DEGs in a specified number of tumor types between RSI groups, ssGSEA and pathway analysis indicates that radiosensitivity is indeed driven by a STAT1-IRF1-CCL4/MIP-1β network that we submit could be exploited to improve the response to radiation.

The RSI was derived from the NCI-60 cell line panel under uniform culture conditions by modeling the relationship between the basal molecular repertoire and clonogenic survival following a clinically relevant radiation dose.^4^ So, how does one explain the dominant ICI signal identified in patient tumors by the RSI, which was derived in cell culture devoid of ICIs? We hypothesize that promiscuous genomic instability^48, 49^ and replication stress^50^ manifest in these cell lines produces danger associated molecular patterns (DAMPS; e.g., cytosolic DNA) and provokes a chronic IFN-based stress response^51, 52^ that was identified in *in vitro* assays of radiosensitivity. The data are consistent with a model where an evolutionary conserved process of ‘viral mimicry’ that is induced by radiation^52^ in tumor cells leads to constitutive IFN signaling via shared STAT1-IRF1 transcriptional stress programs despite the heterogeneous origins of tumors.

Our findings strongly support the notion that the pretreatment TIES (‘hot’-‘altered’-‘cold’)^53^ should be considered when delivering radiation, as the appropriate inflamed state of a tumor^54, 55^ following radiotherapy facilitates tumor kill. Though many clinical trials are testing combinations of immunotherapies with radiation, the optimal dose, fractionation, sequencing and timing of these combinations are unknown. To address this need, we developed and then validated an *in silico* agent-based model that accurately predicts the effects of clinically relevant radiation dosing regimens in tumors harboring varying proportions of interacting effector and suppressor ICIs. Indeed, these simulations predict which TIESs are prone to radiation-induced immune destruction, those that will require priming radiation to shift to a more favorable TIES composition, and those having a resistant TIES where radiation alone will be ineffective. Importantly, these simulations also informed the development of an individual radiation immune sensitivity (iRIS) metric that allows one to more accurately model differences in TIES and personalize radiation dose without compromising efficacy. Specifically, agent-based models informed by each patient-specific pre-treatment ICI composition, RSI and iRIS predicted actual clinical outcomes in NSCLC patients with high accuracy. Critically, this informed model revealed that about half of these lung cancer patients could have potentially de-escalated their radiation dose and still achieved tumor control, and that another 40% required further dose intensification.

The advent of ‘-omics’ analyses has rapidly advanced our understanding of tumor biology and has revealed remarkable heterogeneity of tumor ecosystems and the phenotypes of cells therein. Despite these unequivocal findings, the delivery of radiotherapy, one of the most common therapeutic modalities in oncology, has remained affixed to an imprecise and empiric approach of radiation dose prescription. The era of precision medicine now provides a platform to individualize radiation delivery based on patient-specific tumor attributes coupled with biologically-informed mathematical models.

We provide data that two assumed distinct tumor attributes, radiosensitivity and ICI contexture, are linked. Thus, integrating features of the tumor ICI and radiosensitivity may provide opportunity to individualize radiation dose delivery and assist with deciding whether treatment should be continued, escalated, de-escalated or changed altogether.

## Methods

Methods, including statements of data availability and any associated accession codes and references are available online.

## Acknowledgements

We extend our sincere thanks to the Biostatistics and Informatics Core of H. Lee Moffitt Cancer Center and Research Institute. This work was supported by the NCI Cancer Center Support Grant P30-CA076292 to the Moffitt Cancer Center, by the Cortner-Couch Chair for Cancer Research of the University of the South Florida School of Medicine (J.L.C.), National Institutes of Health R21CA101355 (J.T-R.), De Bartolo Personalized Medicine Institute, and by support from the Florida Breast Cancer Foundation (H.E.) and from the State of Florida to the Florida Academic Cancer Centers Alliance.

## Author Contributions

G.D.G., H.E. and J.T-R. coordinated the study. G.D.G., JC.L.A., E.A.W., S.A.E., J.K.T. and H.E. designed and/or performed experiments. JC.L.A., and H.E performed the *in silico* agent based modeling studies. G.D.G., JC.L.A., E.A.W., S.A.E., J.K.T., H.E. and J.T-R. interpreted data. G.D.G., JC.L.A., K.A.A., L.B.H., S.A.E., J.K.T., J.L.C., J.J.M., H.E. and J.T-R wrote and edited the manuscript. All authors approved the manuscript.

## Extended Data Figure Legends

**Fig. 1.**
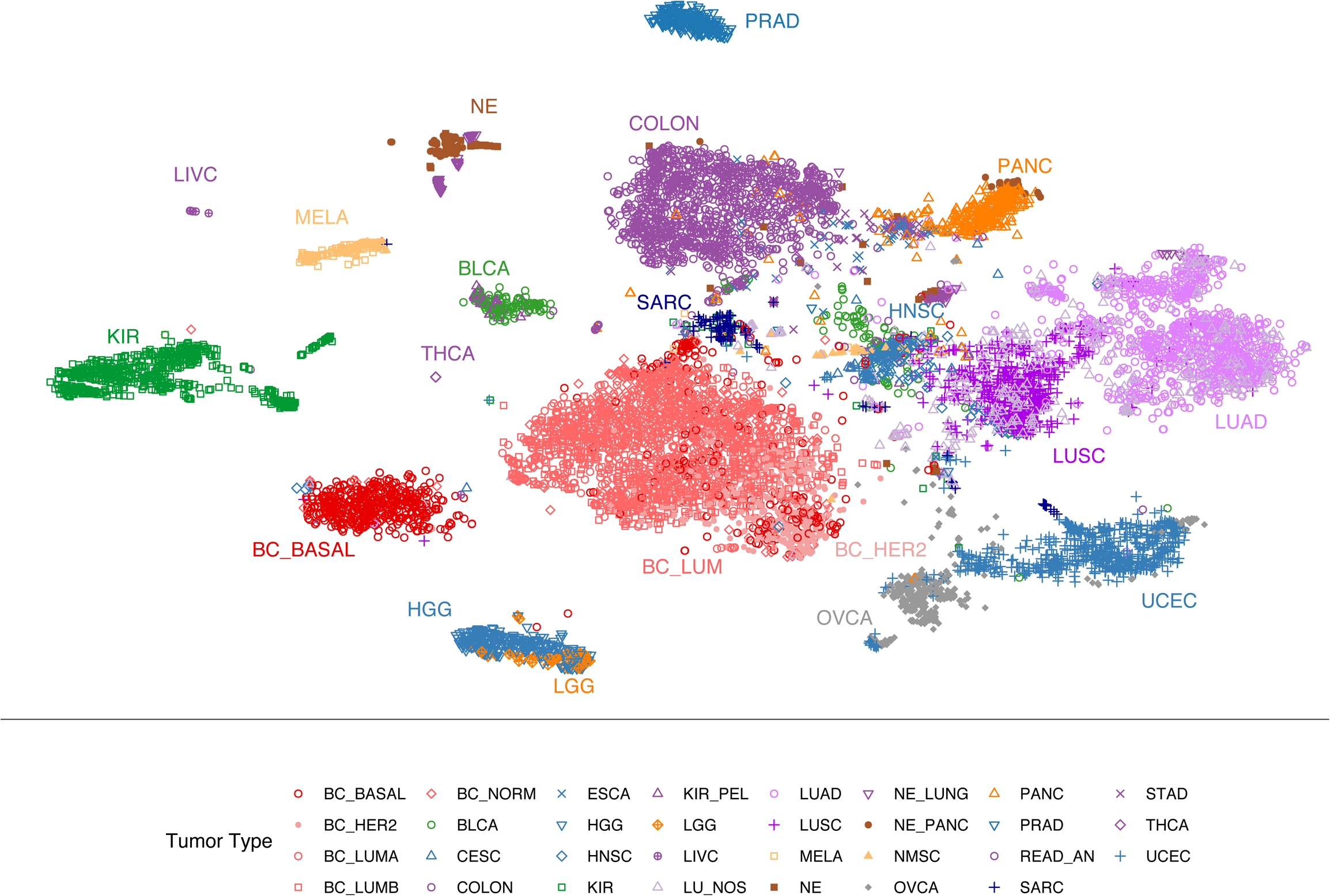
Tumor types cluster by transcriptional programs. Multidimensional reduction analysis of transcription data by the t-distributed stochastic neighbor embedding (t-SNE) method demonstrates distinct clustering of tumor types. t-SNE plot of 10,469 primary tumor samples grouped by tumor type via gene expression profiling.

**Fig. 2.**
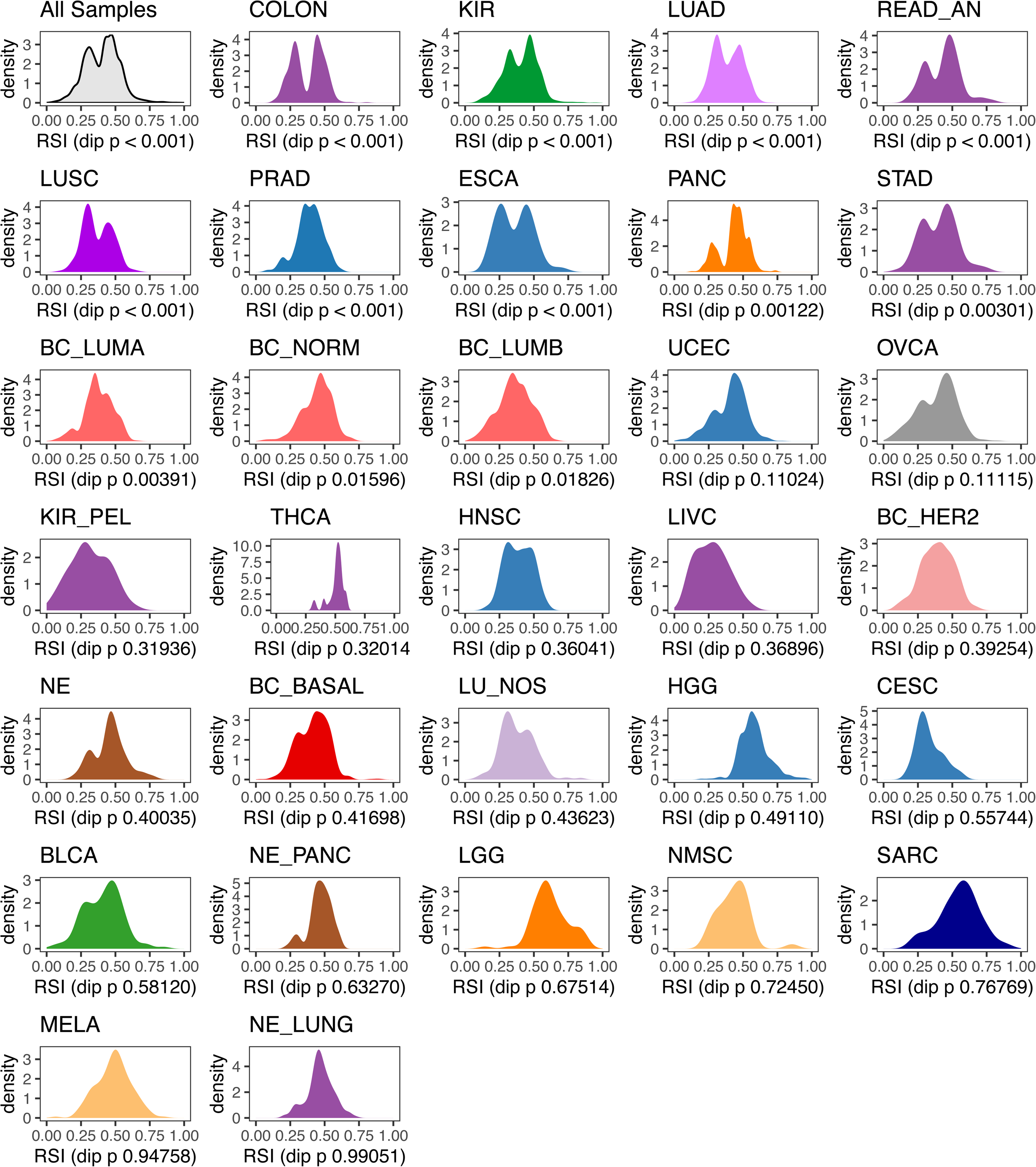
Non-unimodal distributions in RSI. Hartigan’s dip statistic across 31 tumor types to evaluate for unimodal distributions of RSI. Analysis demonstrates that several tumor types do not have a unimodal distribution of RSI values, which suggests biologic heterogeneity.

**Fig. 3.**
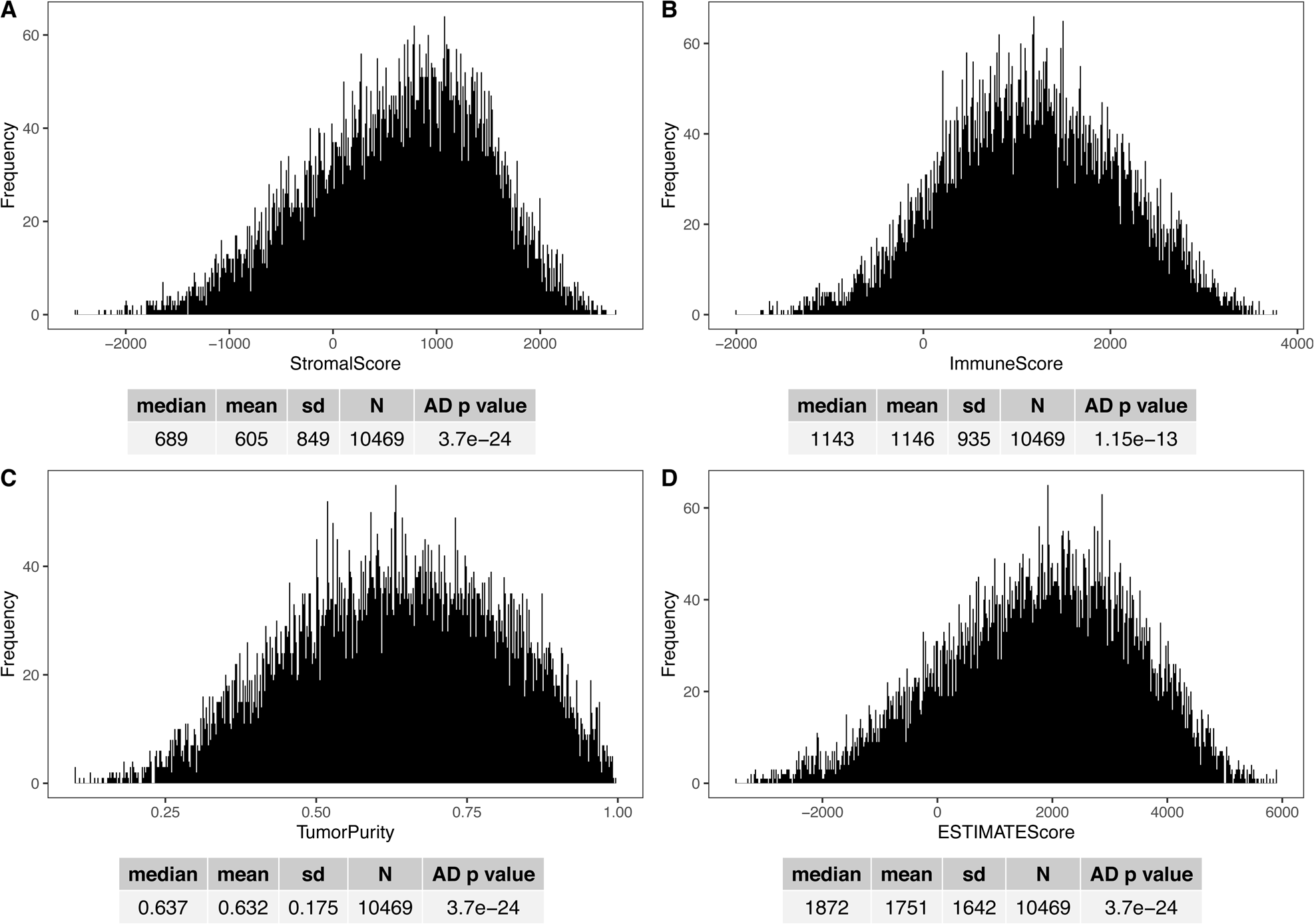
Distributions of ESTIMATE-derived scores. The ESTIMATE algorithm^23^ was used to infer the presence of stromal, immune and malignant cell proportions in 10,469 primary tumor samples in the TCC. Distributions of ESTIMATE-derived stromal and immune scores, as well as tumor purity estimates with associated descriptive statistics.

**Fig. 4.**
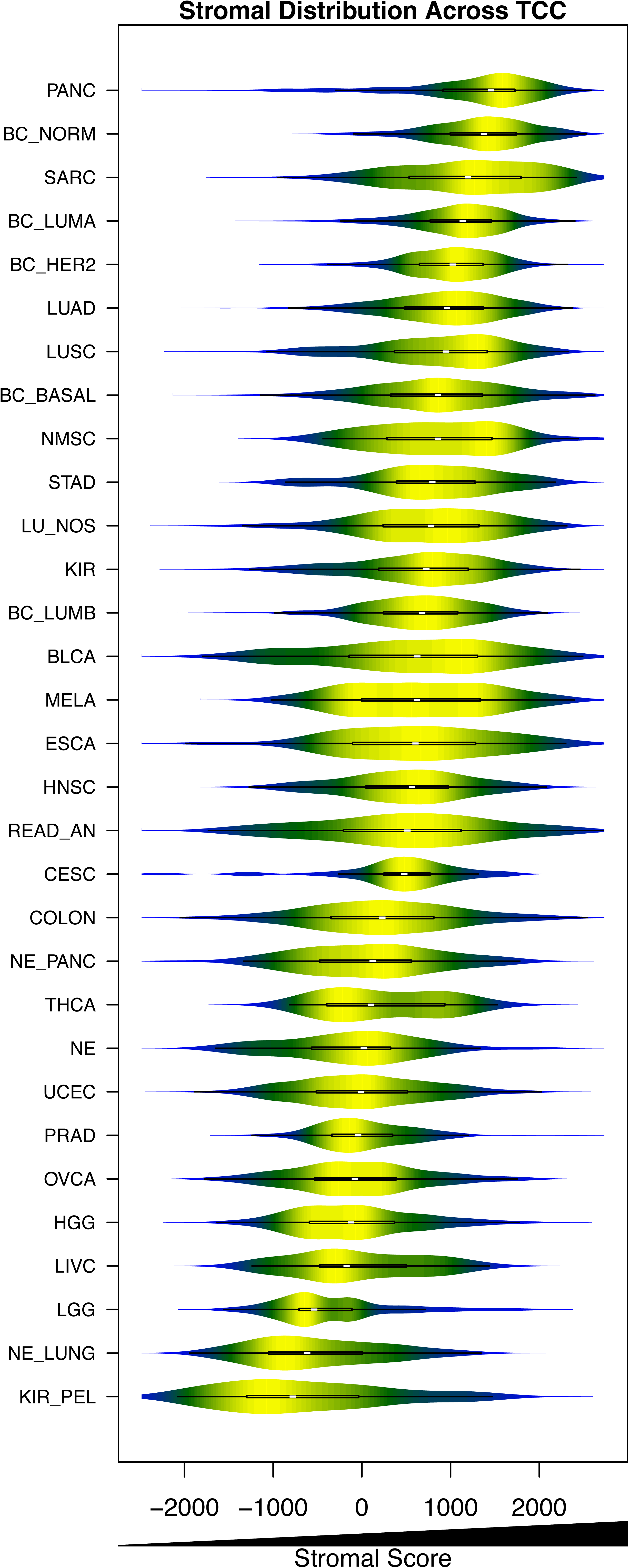
Heterogeneity in stromal cell presence across and within tumor types. Violin plot demonstrating distribution of the ESTIMATE-derived stromal score in 31 tumor types. Demonstrates heterogeneity within and across tumor types for stromal cell presence.

**Fig. 5.**
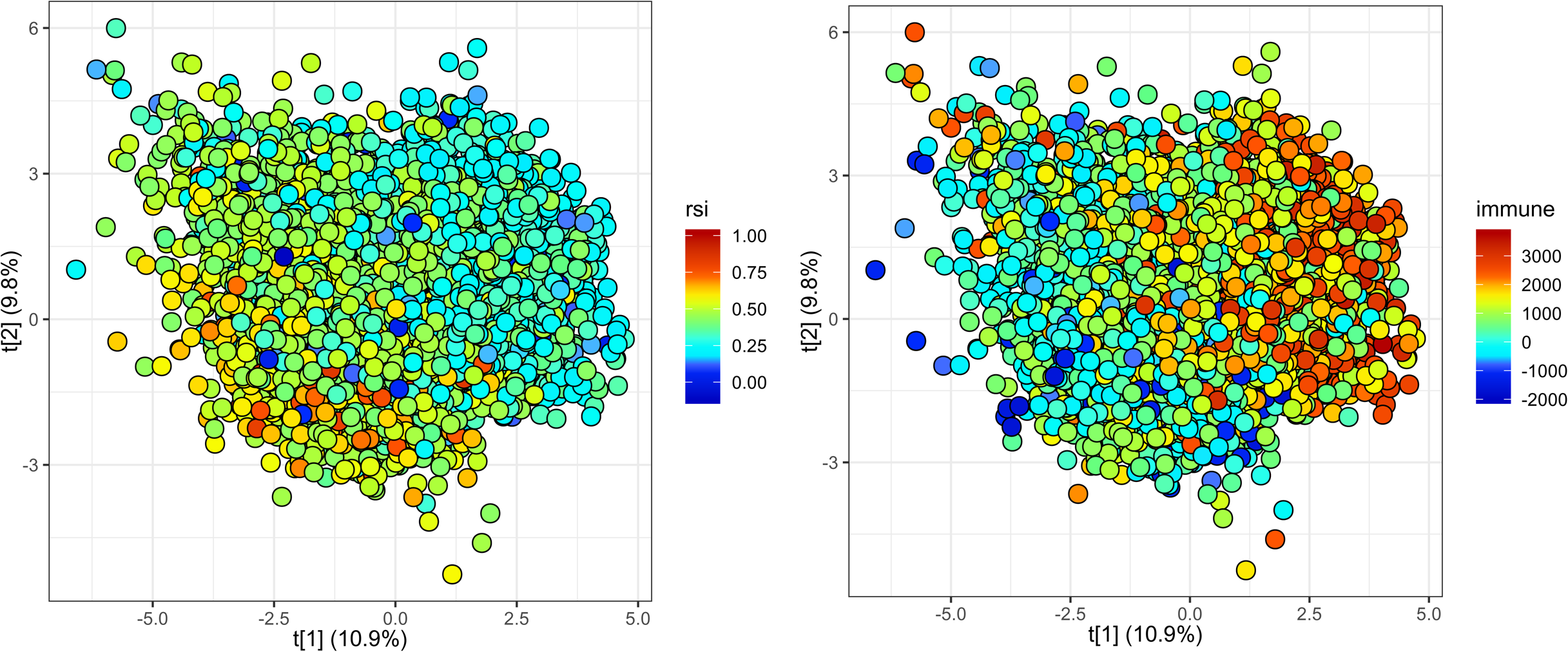
Population level relationship between RSI and ESTIMATE-derived immune score. Score plot of principal component analysis of 10,469 primary tumor samples delineated by continuous RSI (left panel) and ESTIMATE-derived immune score (right panel) values based on gene expression.

**Fig. 6.**
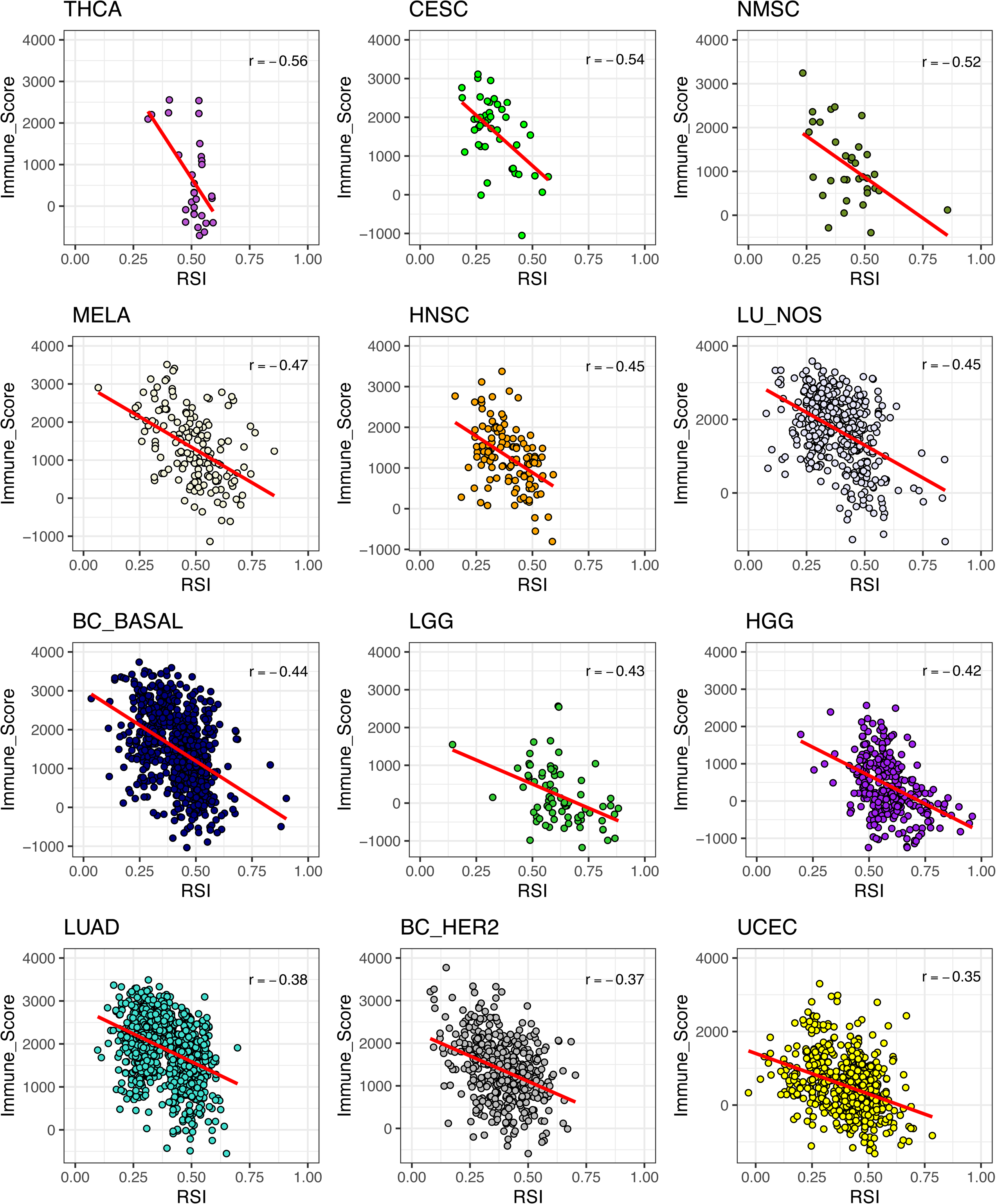
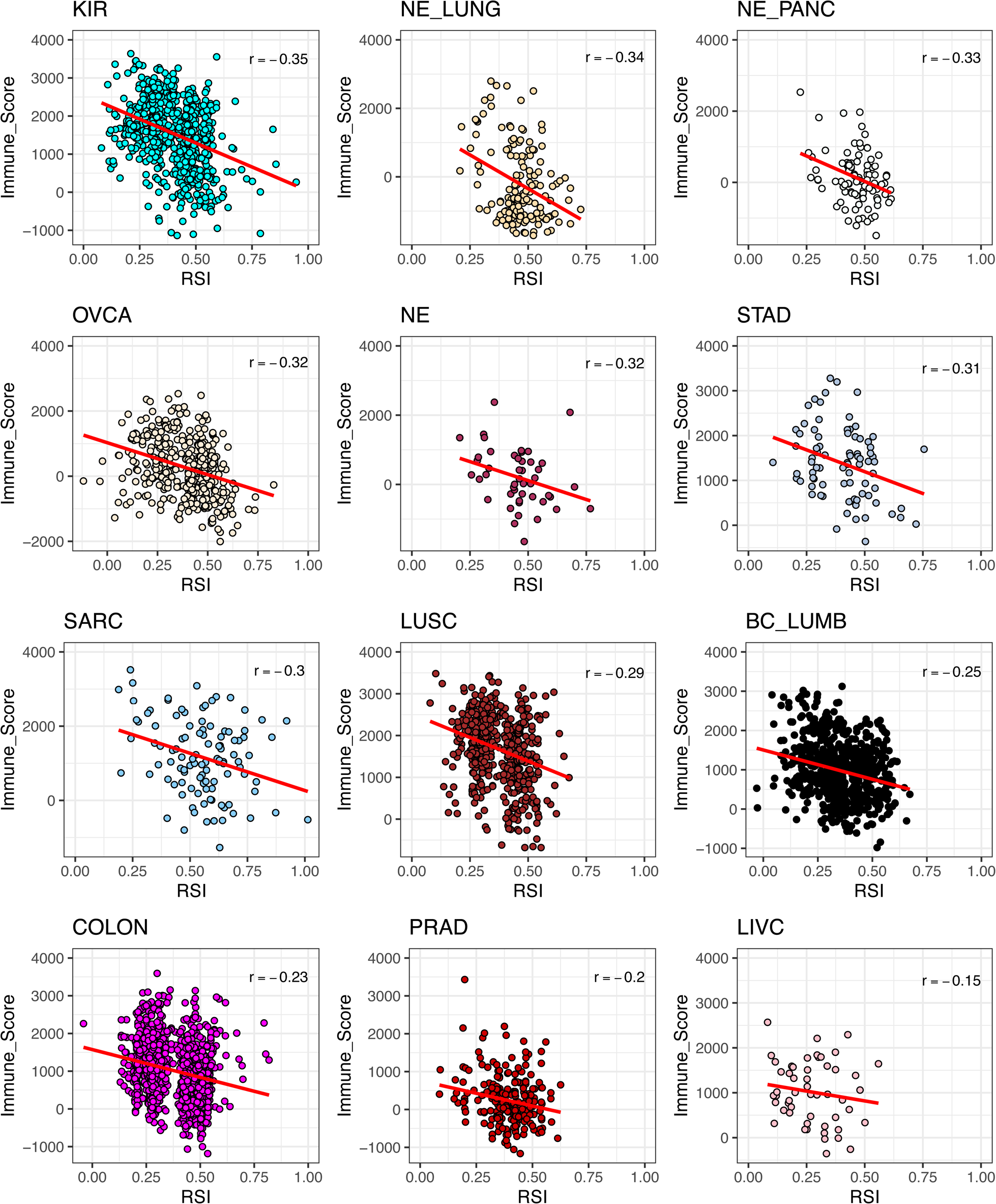
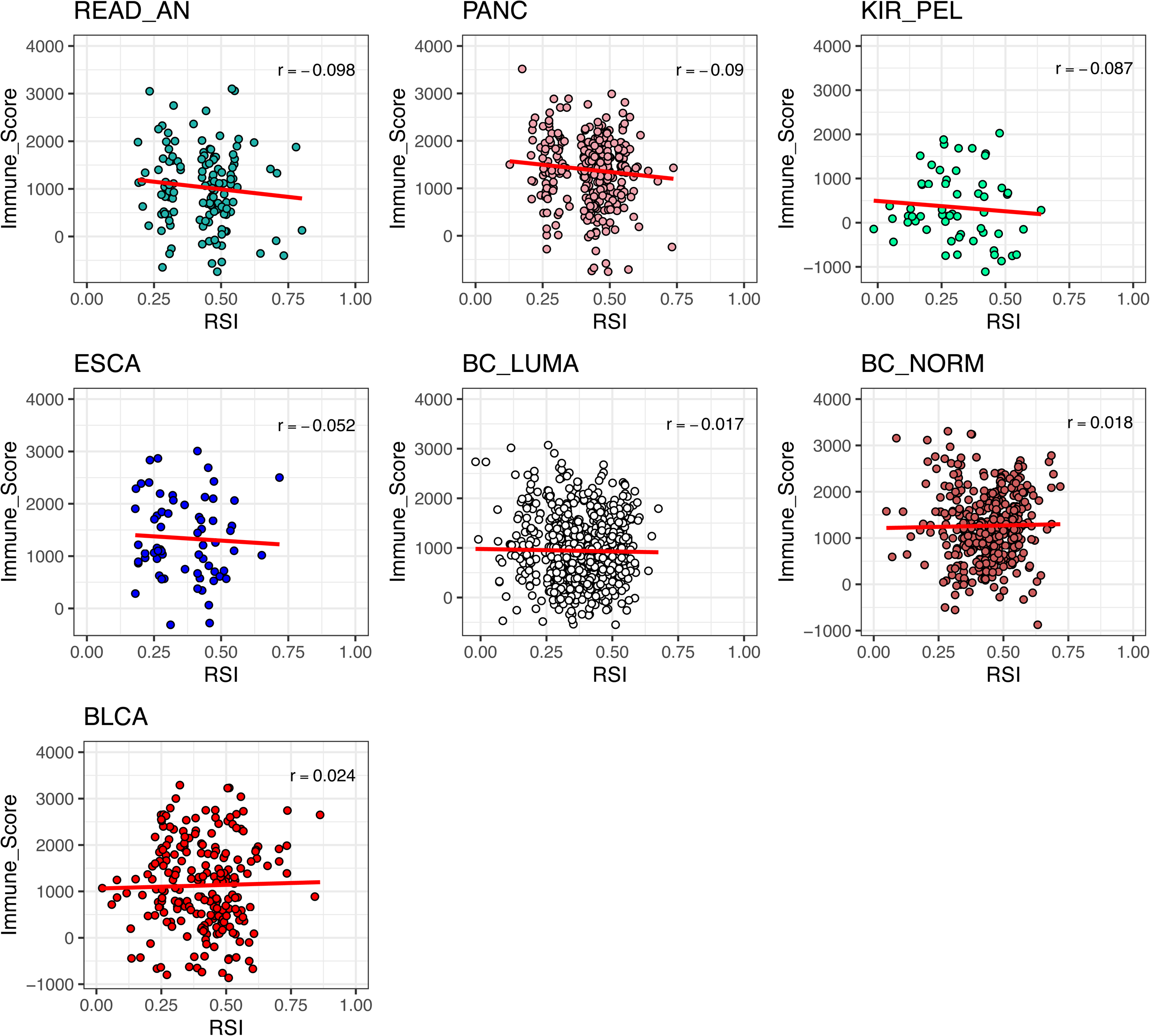
Correlation between RSI and immune score across individual tumor types. Pearson correlation analysis for RSI and immune score among 31 individual tumor types.

**Fig. 7.**
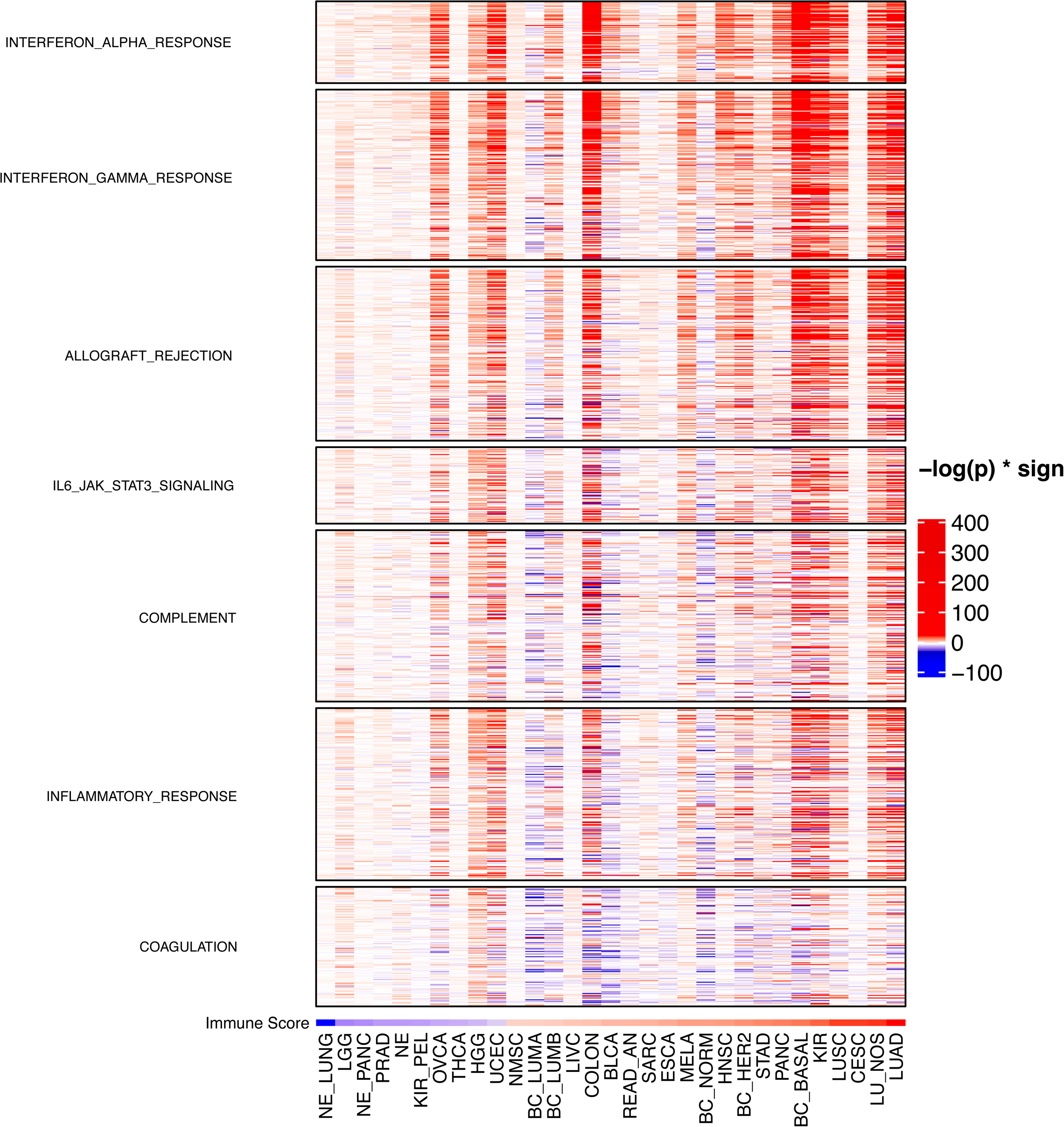
Various immune-related pathways are upregulated in radiosensitive tumors. Heatmap representing the -log p-values of individual gene expression differences of MSigDB hallmark immune pathways between RSI^lo^ and RSI^hi^ tumors. Pathways include those related to IFNα, IFNγ, allograft rejection, interleukin-6-janus kinase-signal transducer and activator of transcription 3, complement, inflammatory response and coagulation signaling.

**Fig. 8.**
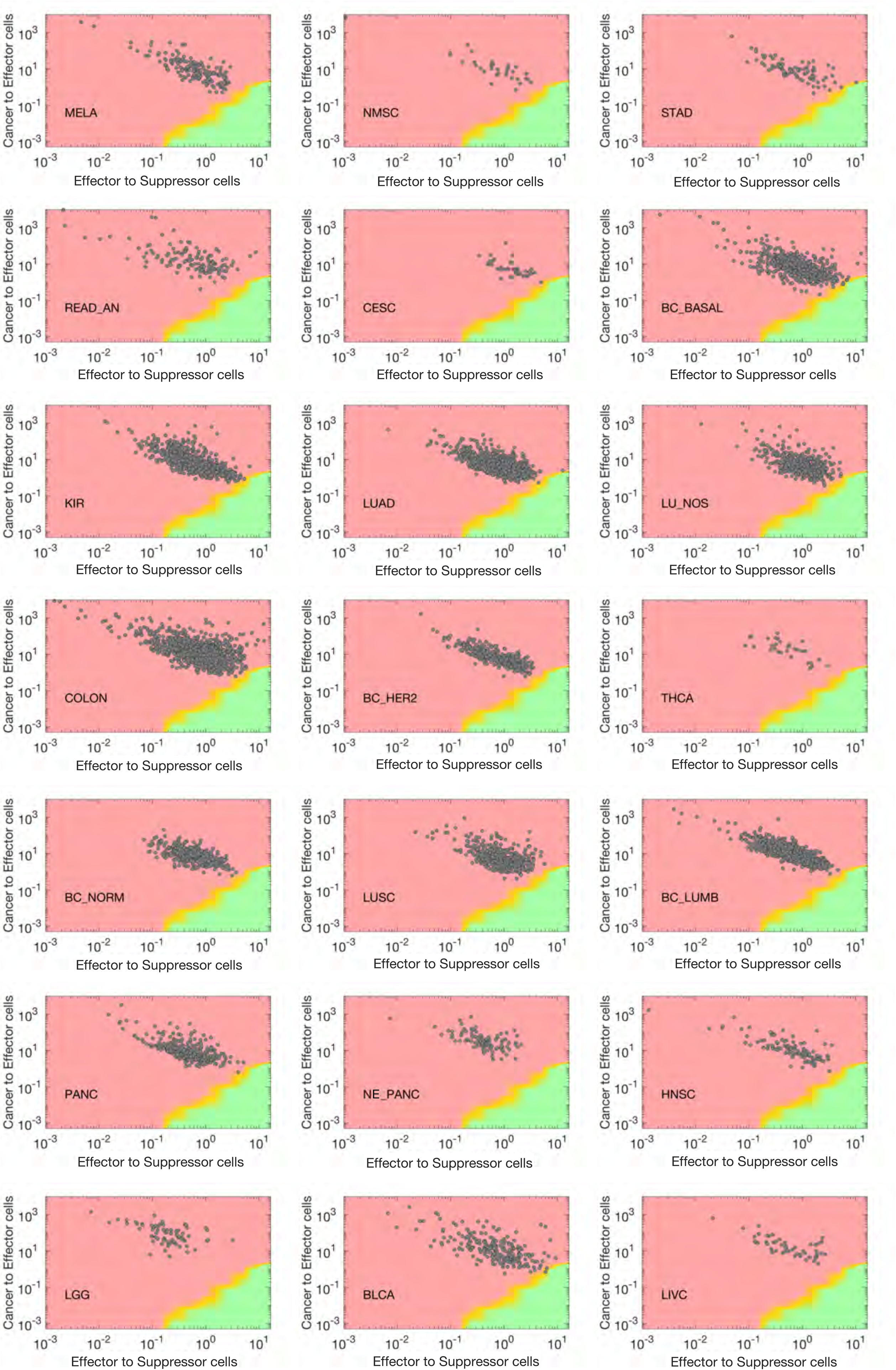
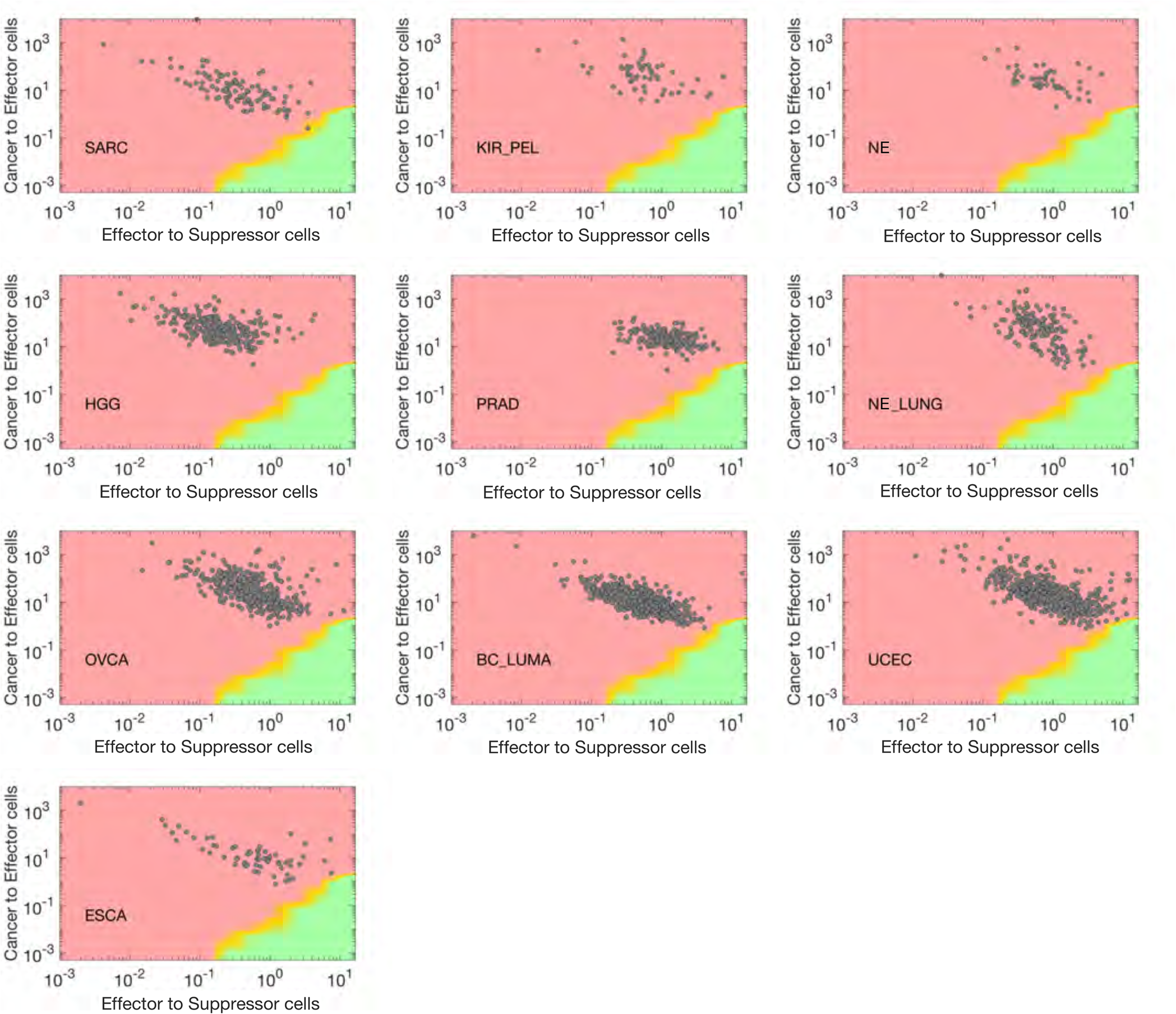
Tumor type specific tumor-immune ecosystem (TIES) maps. TIES maps for 31 tumor types showing probability of immune-mediated tumor elimination (IMTE).

**Fig. 9.**
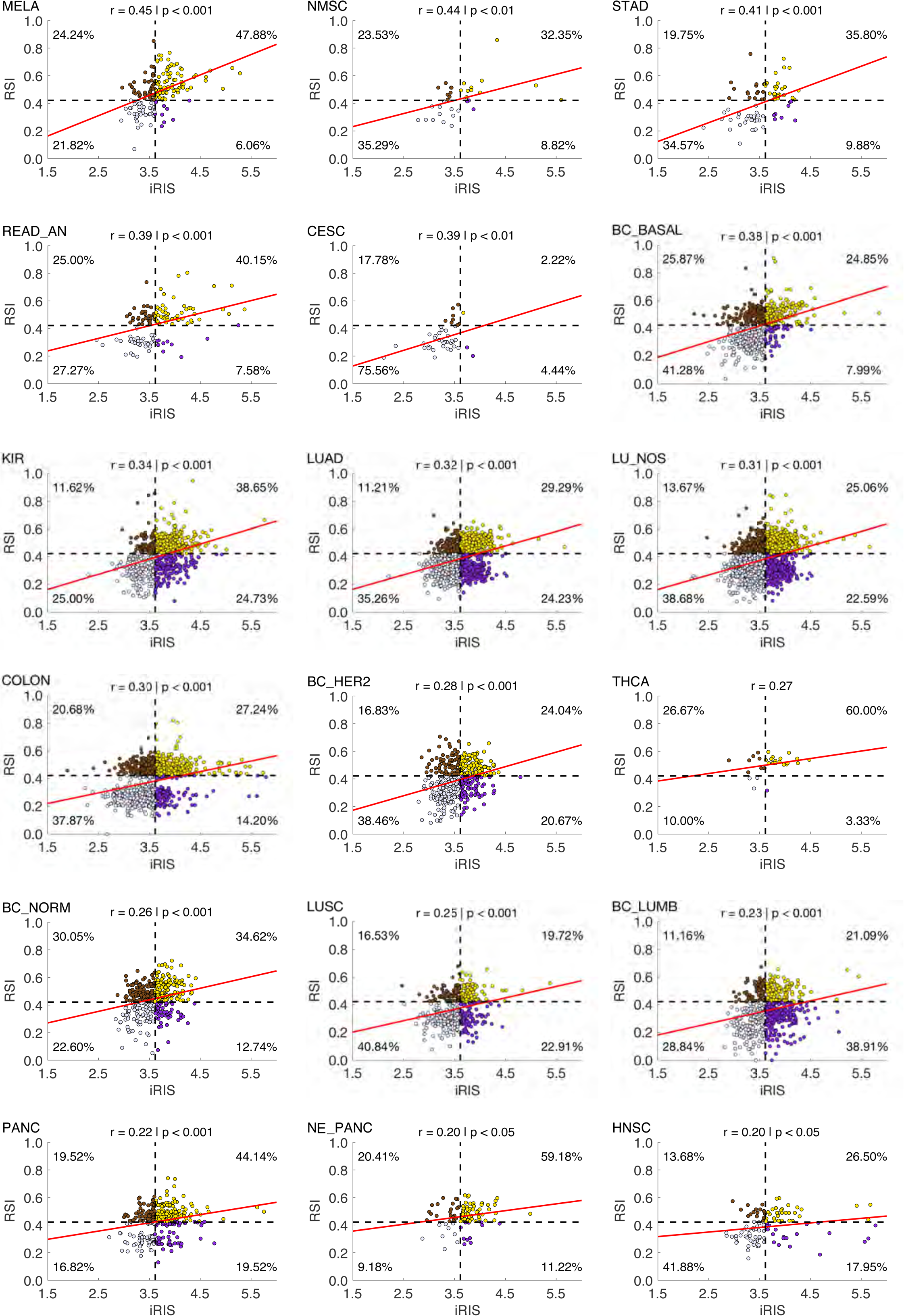
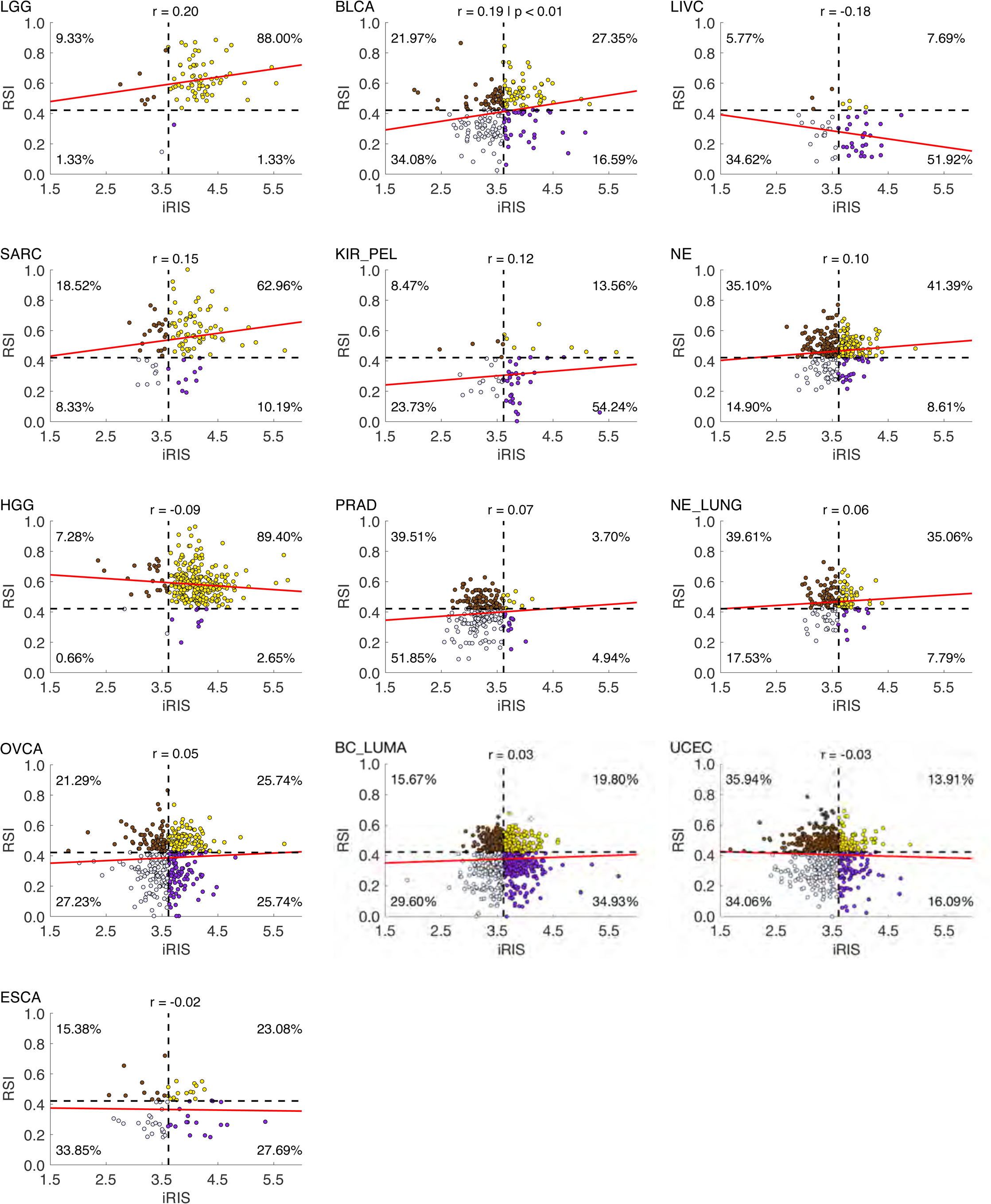
Associations between RSI and iRIS and stratification of tumors by integrating both metrics. Tumors were separated into quadrants by RSI^lo^ and RSI^hi^ (population median RSI value), as well as iRIS^lo^ and iRIS^hi^ (population median iRIS value) into quadrants. Pearson correlations across all 31 tumors types between RSI and iRIS.

**Fig. 10.**
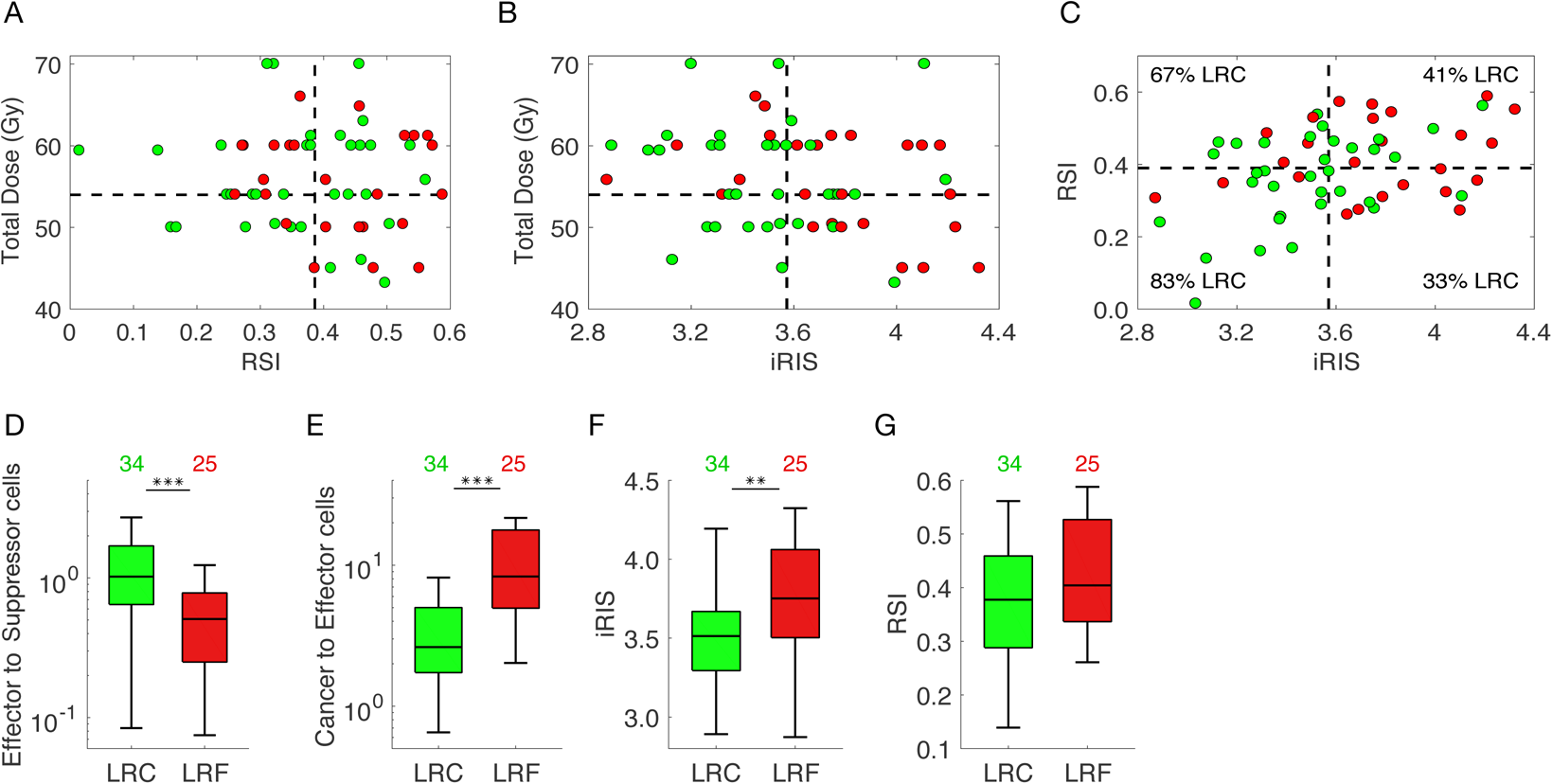
Total radiation dose in NSCLC cohort was not informed by RSI or iRIS, but TIES did predict for enhanced tumor control. **a,b)** RSI and iRIS were not associated with the total radiation dose delivered to the 59 NSCLC patients. **c)** RSI and iRIS were not correlated. **d)** Patient tumors that achieved LRC had **d)** higher effector/suppressor ICI and **e)** lower malignant cell/effector ICI ratios than those with LRF. **e)** Patient tumors that achieved LRC had lower iRIS values compared to those with LRF (*P* <0.01), where **f)** there was no difference in RSI values between patients with LRC or LRF.

## Supplementary Materials

### Methods

#### Patient tumor data set

Patients were consented to the Total Cancer Care^TM^ (TCC) protocol (IRB-approved, Liberty IRB #12.11.0023)^1^. Pathology quality control evaluation of tumors was performed as part of the TCC tissue collection protocol, which includes percent malignant cellularity, necrosis and stromal cell presence. Tumor samples were assayed using the custom Rosetta/Merck HuRSTA_2a520709 Affymetrix gene expression microarray platform (GEO: GPL15048). CEL files were normalized against the median CEL file using IRON^2^, which yields Log_2_ intensity values per probeset. Principle component analysis (PCA) of all samples revealed that the first principle component was highly correlated to RIN value, suggesting a RNA quality difference among samples. A partial least squares (PLS) model was trained upon the fresh frozen samples for which RIN was available and used to re-estimate the RNA quality of all samples. Then the first principle component was removed to correct the signals for RNA quality. 10,469 fresh frozen macrodissected primary tumors were identified for the related analyses.

#### Moffitt non-small cell lung cancer cohort

This cohort was obtained from archived tumors that were resected between 2000 and 2010 from patients in the TCC and Moffitt Cancer Center database. All patients provided written informed consent tissue acquisition, molecular profiling and follow-up. We identified 59 tissue samples, which were surgically resected and pathologically confirmed, American Joint Committee on Cancer version 6, stage IIIA-IIIB tumors. Each patient underwent post-operative radiotherapy. Time to recurrence and overall survival was assessed by determination of the treating physician based on clinical source documentation. Gene expression data was obtained from the TCC.

#### tSNE

Using the 10,469 samples, we first filtered the 60,607 probesets to those having a standard deviation (across the cohort) > 2, yielding 1,399 probesets. The NIPALS algorithm (R/nipals package v0.5) was used to compute the first 50 principal components of the expression data. Rtsne (v0.15) was used with perplexity=100 to generate a two-dimensional mapping from these principal components.

#### Violin Plots

Plots were generated using R (3.6.0) using an in-house custom code (https://github.com/steveneschrich/RSI_Immune_Paper). The violin plots represent the smoothed density with nrd0 bandwidth kernel.

#### Estimate

The ESTIMATE algorithm^3^ was used to evaluate stromal and immune cell fractions from gene expression of the primary tumor (Estimate R package v1.0.13). Normalized gene expression probeset identifiers were mapped to Entrez GeneIDs. As defined in the Estimate package, the data were filtered to only the common genes. A single probeset was then selected per gene by choosing the probeset having the highest median expression across all tumors. This resulting dataset was used in the estimate function to produce stromal and immune scores for each cohort.

#### CIBERSORT

The HuRSTA array was reduced to the LM22 signature genes as defined by CIBERSORT^4^ by choosing a representative probeset that detects the gene and has the highest median expression among matching probesets. The CIBERSORT web tool (https://cibersort.stanford.edu/index.php) was accessed on 2017-05-19 to generate signature scores (using quantile normalization).

#### Immune Cell Infiltrate (ICI) Normalized Content Abundance

ICI composition proportions from CIBERSORT^4^ were extracted in relative mode, where the proportions are relative to the total immune cell fraction of the tumor. To normalize the content across tumors, we scaled the ESTIMATE^3^ immune scores such that the lowest immune score was 0 (rather than negative) and analyzed multiple ICI fractions by this adjusted immune score, to yield the Normalized Content Abundance (NCA).

#### ICI PCA Plots

Relative ICI composition proportions (CIBERSORT estimates) were used to compute principal components across the entire TCC cohort using prcomp (R3.6.0) with scaling. The loadings and scores were visualized using R packages ggplot2 (3.2.1), ggrepel (0.8.1), and ggpubr (v0.2.3).

#### ICI Enrichment

RSI was dichotomized into RSI_HIGH and RSI_LOW by tumor subtype using the median RSI for that type. Each TIL (and ESTIMATE score) was likewise dichotomized using the same approach into ICI_HIGH and ICI_LOW. A Fisher exact test was performed for each comparison of dichotomized RSI and ICI to determine the independence of the variables. The q value package (2.16.0) was used to compute FDR q values to correct for multiple testing^5^ (* see additional references). The ICIs were classified as composing adaptive and innate cell types based on current literature and PCA. The enrichment heatmap was generated using ComplexHeatmap (2.0.0)^6^. Each cell represents the proportion of samples with ICI_HIGH in either RSI_LO (top row) or RSI_HI (bottom row). The percentage of disease sites with a significant q value (q<0.05) of differences between ICI_HIGH in RSI_LOW vs. RSI_HIGH is noted.

* Storey JD. The Positive False Discovery Rate: A Bayesian interpretation and the Q-value. *Annals of Statistics* **31**, 2013-2035 (2003).
* Storey, JD, Taylor JE, and Siegmund D. Strong control, conservative point estimation and simultaneous conservative consistency of False Discovery Rates: A unified approach. *J Royal Statistical Society. Series B (Statistical Methodology)* **66**, 187-205 (2004).

#### Calculation of RSI

RSI was calculated for each sample using the within-sample rank coefficients as described^7, 8^. The following 10 probesets, corresponding to the originally defined HG-U133plus2 probesets, were used: merck-NM_007313_s_at (ABL1), merck-NM_000044_a_at (AR), merck-NM_004964_at (HDAC1), merck-NM_002198_at (IRF1), merck-NM_002228_at (JUN), merck-BQ646444_a_at (PAK2, CDK1), merck2-X06318_at (PRKCB), merck2-BC069248_at (RELA), merck-NM_139266_at (STAT1), merck-NM_001005782_s_at (SUMO1).

#### ssGSEA

Single-sample GSEA^9–11^ was used to evaluate the MSigDB Hallmarks Genesets^12–14^. The GSVA^15^ package (R package GSVA, v1.32.0) was used to calculate ssGSEA scores. ssGSEA scores were combined by tumor type using the median score.

#### RSI^lo^ *vs.* RSI^hi^

For probesets that were differentially expressed (DE) between RSI^lo^ and RSI^hi^ within each tumor type, samples were ranked by RSI score and divided into RSI^lo^ and RSI^hi^ groups at the median RSI value. A probeset was determined to be DE within a tumor type if the following criteria were met: (i) probeset must not be annotated as antisense; (ii) one of the two group averages must be > 5; (iii) |log2 ratio| ≥ ∼0.585 (1.5-fold); (iv) two-tailed t-test ≤ 0.01; and (v) Hellinger distance ≥ 0.25.

#### Metacore pathway

Probesets that were identified as DE in ≥ 6 tumor types and that had no conflicting direction of change (197 probesets, 146 genes) were used as seeds for MetaCore (Clarivate Analytics) literature network generation. Networks were generated using the Build Network-Shortest Paths method with the following non-default settings: (i) do not show disconnected seed nodes; (ii) do not show shortest path edges only; and (iii) use only Phosphorylation, Dephosphorylation, and Transcription Regulation interaction types of known Activation / Inhibition status. The resulting interaction network was exported and further filtered to contain only nodes and edges consistent with the observed RSI^lo^ vs. RSI^hi^ directions of change, which were then visualized using Cytoscape v3.3.^16^

#### DNA sequencing and analysis

Sequencing analysis are described in detail as previous^17^. Briefly, coding regions of 1,321 genes were targeted with SureSelect (Agilent Technologies, Santa Clara, CA) custom capture and sequenced on GAIIx sequencing instruments (Illumina, Inc., San Diego, CA) in paired-end 90bp configuration. Sequence reads were aligned to the reference human genome with the Burrows-Wheeler Aligner (BWA)^18^, and duplicate identification, insertion/deletion realignment, quality score recalibration, and variant identification were performed with PICARD (http://picard.sourceforge.net/) and the Genome Analysis ToolKit (GATK)^19^. Genotypes were determined with GATK UnifiedGenotyper and refined using Variant Quality Score Recalibration. Variants were filtered based on quality (highest sensitivity tranche were excluded) and population frequency (variant observed in 1000 Genomes Project or NHLBI Exome Sequencing Project or at >1% in 238 normal samples from the TCC cohort were excluded). Sequence variants were annotated to determine genic context (ie, non-synonymous, splicing) using ANNOVAR^20^.

#### Statistics

Statistical analysis was done using R 3.6.0. The RSI distribution across all tumor types and for each specific tumor type was tested for being unimodal using Hartigan’s dip test (R package diptest, v0.75-7) and visualized using ggplot2 (3.2.1) and ggpubr (v0.2.3). Distributions of ESTIMATE scores (immune, stromal, tumor purity, ESTIMATE) were tested for normality using the Anderson-Darling test for normality (R package nortest, v1.0-4) and visualized using ggplot2 (v3.2.1) and ggpubr (v0.2.3). RSI and Immune score were compared using Pearson correlation (r) and visualized using ggpubr (v0.2.3). RSI and all ESTIMATE scores (stromal, ESTIMATE, immune and tumor purity) were compared using GGally (v1.4.0).

## Mathematical Modeling Methods

### Agent-based model infrastructure

We developed a three-dimensional (3D) multiscale agent-based model that simulates the interactions of cancer cells with anti-tumor immune effector T-cells and immune-inhibitory suppressor cells. Each cell is considered as an individual agent, and their behavior at any time is determined by a stochastic decision-making process based on biological-driven mechanistic rules (see refs. 1-4 for similar modeling approaches). Immune cells are simulated on a 3D regular lattice divided into 200 x 200 x 200 nodes representing a volume of 65mm^3^, with a lattice constant of 20μm (average cell diameter, ref. 3). Cancer cells occupy an irregular 3D lattice that is initialized by randomly placing one lattice node inside each cube of the regular lattice for immune cells. These interconnected lattices are then used to simulate the infiltration of immune cells into the tumor bulk, thus avoiding contact inhibition between immune and tumor cells. A Moore neighborhood is considered for nodes in the same lattice; i.e., 26 orthogonally and diagonally adjacent lattice sites, where each node has eight neighbor nodes in the opposing lattice given by the nodes surrounding it.

Model simulations are initiated with a single cancer cell at the center of the 3D simulation domain. At each time-step (*Δt = 1h*), trafficking, motility, cytotoxic function and suppressing activity of immune cells, as well as tumor cell processes are executed through an iterative procedure (**Supplementary Fig. 1**). Every time-step the state and dynamics of all cancer and immune cells are updated in random order to avoid order biases. When the tumor reaches a population of 10^5^ cancer cells, the number of cancer cells, immune effectors and suppressor cells define the pre-radiation tumor-immune ecosystem (TIES). The direct dose-dependent cytotoxic effect of radiotherapy (RT) on all participating cell types is simulated using the established Linear-Quadratic (LQ) model^5–7^. Model simulations continue after RT until either the immune system eradicates remaining cancer cells, or recurrent tumors reach the pre-treatment number of 10^5^ tumor cells.

### Cancer cell dynamics

Cancer cells proliferate, migrate or undergo programmed cell death (apoptosis) at pre-defined intrinsic probabilities modulated by the local microenvironment (**Supplementary Fig. 1A**). Proliferation and migration attempts are only successfully executed for cells residing at a lattice node with at least one vacant adjacent node. During mitosis, the new cancer cell is placed on a randomly selected free neighbor node. Similarly, cells migrate into adjacent vacant lattice nodes with equal probability to simulate random walks and cell diffusion (Brownian motion). The ability of the tumor cell to proliferate and move is temporarily lost due to contact inhibition, and this quiescent state is abandoned as soon as at least one neighbor lattice site becomes free. Cancer cells that undergo programmed cell death with specified probability are removed from the simulation domain instantaneously. For demonstration purpose, we assume commonly used generic cancer cell parameters, including a cell cycle duration of 35 hr (which results in a proliferation probability *p_p_* = 0.02.31 x 10^-2^ per time-step (*Δt = 1h*)^8^), and apoptotic and migration probabilities as *p_a_* = 4.17 x 10^-3^ (approximately every 10 days^9^) and *p_m_* = 4.17 x 10^-1^ (about cell widths per day^10^).

**Supplementary Fig. 1.**
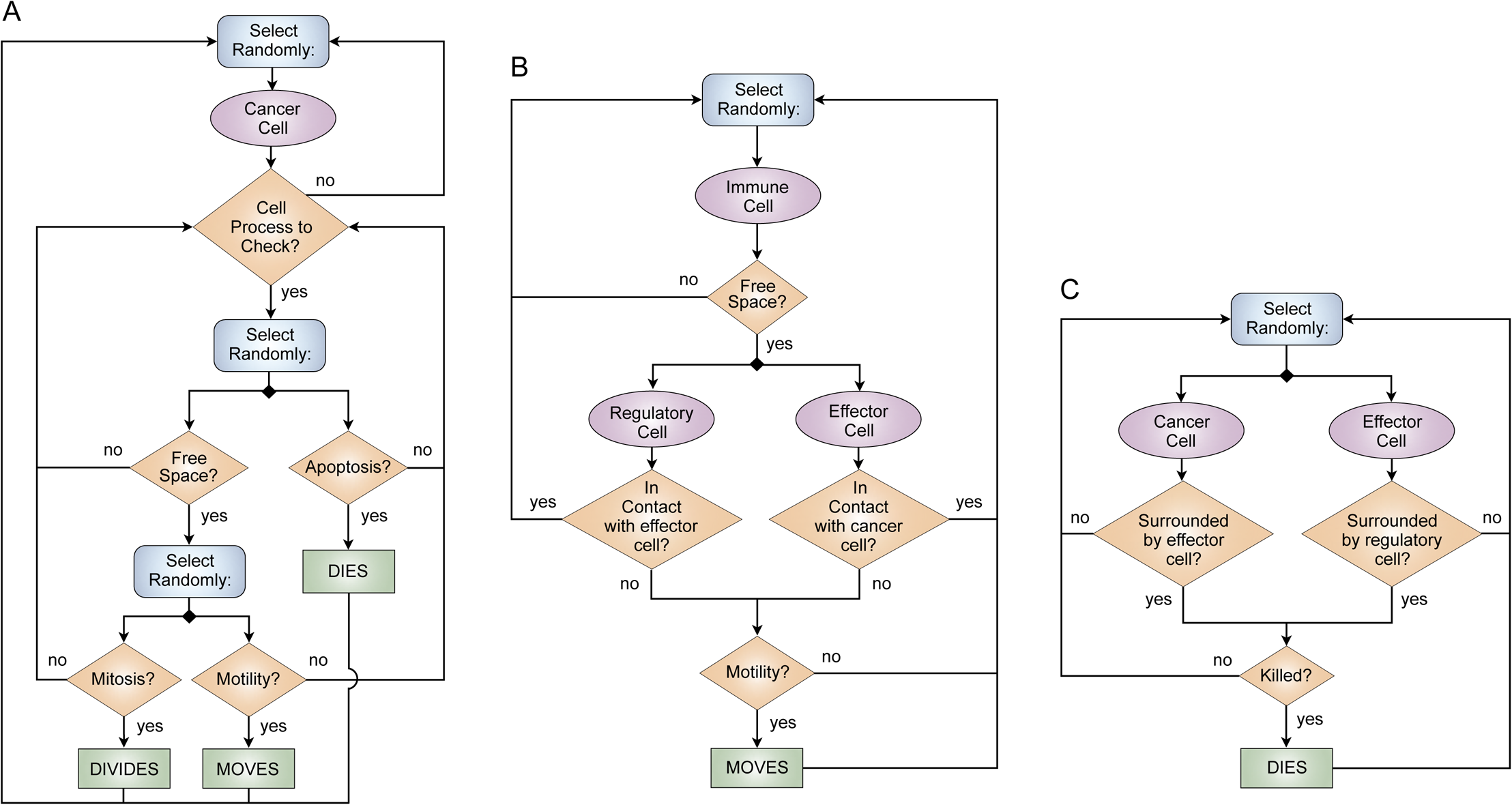
Cell decision iterative flow diagrams. A, Cancer cell decision-making process. B, Immune cells motility process. C, Cell-cell interactions process.

### Immune cell infiltration into the tumor microenvironment

Immune cells arrive in the tumor microenvironment after extravasation from the vasculature^11–13^. Blood vessels are assumed to be homogeneously distributed throughout the simulation domain, and immune cells enter the simulation at randomly selected free lattice nodes in the domain. At each simulation time-step t_i_, the number of effector and suppressor cells that are recruited in response to tumor burden is given as:

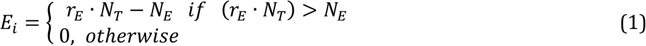

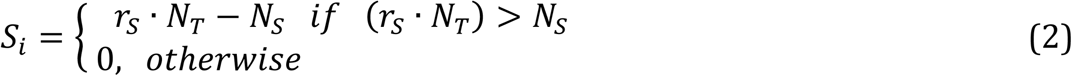

where *N_T_*, *N_E_* and *N_S_* are the amount of tumor cells (T), effector cells (E) and suppressor cells (S) in the system at time t_i_. By considering *r_E_ = N_E_ / N_T_* and *r_S_ = N_S_ / N_T_*, pre-defined ratios of effector-to-tumor cells and effector-to-suppressor cells can be simulated. For patient-specific simulations, the surgical tissue-derived number of cancer, effector and suppressor cells initiate the RT model *(C_0_, E_0_ S_0_),*.

### Immune cell motility

Immune cells migrate and infiltrate tumors via chemotaxis along cytokines and chemokine gradients that are secreted by cancer cells or other immune cells^14^. The mean velocity of immune cells in non-lymphoid tissues have been estimated at 4–10 μm min^-1^, with a peak velocity as high as 25 μm min^-1^ in the lymph nodes^15–17^. Here, we set the mean immune cell velocity to 10 μm min^-1^, which results in a lattice move probability of *h_r_ = 0.37* per min^-1^. The waiting time for immune cell movement follows an exponential probability distribution. Immune cell motility is updated 60 times (every minute) per simulation time-step (*Δt = 1h*). Both effector and suppressor cells move towards the tumor center of mass subject to at least one vacant lattice node in their immediate neighborhood and direction (**Supplementary Fig. 1B**). Immune effector and suppressor cells only move while not performing cytotoxic or suppressing functions, i.e., when not in contact with cancer cells or effector cells, respectively.

Immune cells select free neighbor lattice nodes with a certain probability depending on their distance to the center mass of the tumor. The transition probability for the directed movement of an immune cell from a lattice site *r* to a free neighbor *r’* is given by:

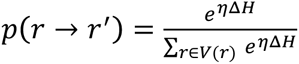

where *V(r)* is the set of free neighbor lattice nodes of *r*, *ΔH = H(r’) -H(r)*, and *H(j)* is the distance of a lattice site *j* to the tumor center of mass 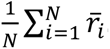. Here the vector *r̄_i_* denotes the spatial coordinates of the lattice node *r_i_* occupied cell and *N* is the total amount of tumor cells. A random number is generated to select a free neighbor lattice site for movement, which is determined from the set of probabilities into which the generated number falls in. We fixed *η* = 0.25, which results in a non-deterministic biased random walk.

### Cell-cell interactions

Accumulating evidence demonstrates cancer cells secrete a wide array of different chemokines and chemotactic cytokines that recruit pro- and anti-tumor immune cell subsets to the tumor microenvironment^14^. While cytotoxic effector T cells have the potential to kill cancer cells by inducing different forms of cell death, suppressor T cells suppress anti-tumor immunity by inhibiting the cytotoxic responses of effector T-cells^18^. Accordingly, effector T cells can kill cancer cells by direct contact with probability *p_E_* = 0.03 per time-step, and suppressor T cells can suppress effector-dependent responses by repressing effector T cells when they are in contact at a probability *p_R_* = 0.01 per time-step (**Supplementary Fig. 1C**). Tumor cell killing and effector T cell inactivation probabilities increase proportionally with the number of immune cells of the same types in the immediate cancer cell neighborhood. The target cell is removed from the simulation domain.

### Effect of radiation on tumor and immune cells

The cytotoxic effect of radiotherapy on cancer cells was simulated by using the standard Linear-Quadratic (LQ) model^5–7^. The radiation dose *D* (Gy) dependent surviving fraction (SF) is given by:

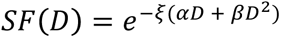

where *α* (Gy^-1^) and *β* ^-^(Gy^-2^) are cell type-specific radiosensitivity parameters. Growth arrested cells are approximately three-times more resistant to radiation than normoxic cycling cells^19^; we set *ξ* = 1 and *ξ* = 1/3 to respectively scale the radiosensitivity of proliferative and quiescent tumor cells as previously discussed^2^.

As *in silico* tumors are about six orders of magnitude smaller than clinically apparent tumors, simulating seven weeks of radiation will overestimate tumor control. With generic cancer radiosensitivity of *α* = 0.3 Gy^-1^ and *β* = 0.03 Gy^-1^, we have set SF (2 Gy x 30) ≅ 4e-10. With SF (2 Gy x 10) ≅ 7e-4, each RT dose *in silico* is estimated to equate three actual radiotherapy doses. For RT simulations for an individual NSCLC patient *k*, we set SF*_k_* (2 Gy) = RSI*_k_*.

Immunosuppressive cells are intrinsically more resistant to radiation than cytotoxic effector T cells^20, 21^, in line with estimates of radiation-induced lymphocyte death after increasing radiation doses *in vitro*^22^. From these dose-response curves we derive SF_S_(1.8Gy) = 0.81 and SF_S_(2.0Gy) = 0.79 for suppressor immune cells, and SF_E_(1.8Gy) = 0.63 and SF_E_(2.0Gy) = 0.61 for immune effector cells.

### Radiation-induced recruitment of immune cells

Radiation induces immunogenic cell death, and this releases immune-stimulating signals and enhances T cell infiltration into the TME^20, 23–25^. Radiation-induced tumor cell death occurs during the first hours after irradiation^26^ and is marked by increased infiltration and recruitment of T lymphocytes^27^. Radiation-induced effector and suppressor cells recruited at time *t_i_* is simulated by:

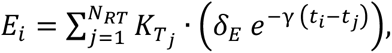

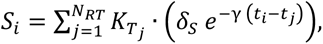

where *N_RT_*, is the number of radiation fractions, and K_Tj_ is the amount of cancer cells eradicated by radiation at fraction *j* delivered at time *t_j_*. The parameters δ_E_ and δ_S_ describe radiation-induced recruitment rates of effector and suppressor cells, and γ is the decay of radiation-induced immune stimulation. Parameter values δ_E_ = 0.05 h^-1^, δ_R_ = 0.01 h^-1^ and γ = 0.05 were used for all model simulations.

### Individual Radiation Immune Sensitivity (iRIS) score

The TIES composition prior to therapy combines cancer-to-effector cell and effector-to-suppressor cell ratios with the absolute number of suppressor cells. Effector (anti-tumor) ICI was represented by CD8 T, activated CD4 memory T, activated NK, and M1-polarized macrophage cells. Suppressor (pro-tumor) ICI was represented by regulatory T, M2-polarized macrophage and neutrophil cells. These ICI type were chosen in line with previous literature^28^ and clean signal on PCA. Including all the ICI populations following PCA stratification yields comparable results, but lower significance due to ambitious stratification. To correlate TIES composition with radiation responses the individual radiation immune sensitivity (iRIS) score is defined as:

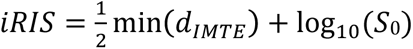

Where

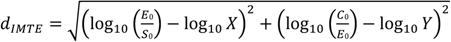

is the distance of pre-radiation TIES to the immune mediated tumor elimination (IMTE) region, and E_0_, S_0_ and C_0_ are the number of effector, suppressor and cancer cells, respectively. The X and Y vectors are the coordinates of all simulated points in the IMTE region in the immune contexture matrix.

